# Immunoglobulin heavy constant gamma gene evolution is modulated by both the divergent and birth-and-death evolutionary models

**DOI:** 10.1101/2021.08.12.456010

**Authors:** Diego Garzón-Ospina, Sindy P. Buitrago

**Affiliations:** PGAME - Population Genetics And Molecular Evolution, Fundación Scient, Tunja, Boyacá, Colombia; GEO, School of Biological Sciences, Universidad Pedagógica y Tecnológica de Colombia - UPTC, Tunja, Boyacá, Colombia; GEBIMOL, School of Biological Sciences, Universidad Pedagógica y Tecnológica de Colombia - UPTC, Tunja, Boyacá, Colombia

**Keywords:** IgG, Primates, Immune system, Multigene family, Birth-death evolutionary model, Divergent evolutionary model

## Abstract

Immunoglobulin G (IgG) is one of the five antibody classes produced in mammals as part of the humoral responses. This high-affinity antibody produced late in a primary immune response is responsible for protecting the organisms from infection. This protein’s heavy chain constant region is encoded by the Ig gamma gene (*Igγ*). In Humans, IgG has evolved into four subclasses with specialized effector functions. However, in *Platyrrhini*, IgG has been reported to be encoded by a single-copy gene. Here, we analyzed data from 38 primate genome sequences to identify *Igγ* genes and describe the evolution of this immunoglobulin in this group. *Igγ* belongs to a multigene family that evolves by the birth-death evolutionary model in primates. Whereas *Strepsirrhini* and *Platyrrhini* have a single-copy gene, in *Catarrhini* species it has expanded to having several paralogs in their genomes; some have been deleted or have become pseudogenes. Furthermore, episodic positive selection might promote a specialized species-specific IgG effector function. A hypothesis for the *Igγ* evolution has been proposed, suggesting that IgG has evolved to reach an optimal number of copies per genome to adapt their humoral immune responses to different environmental conditions.

**Graphical abstract:** 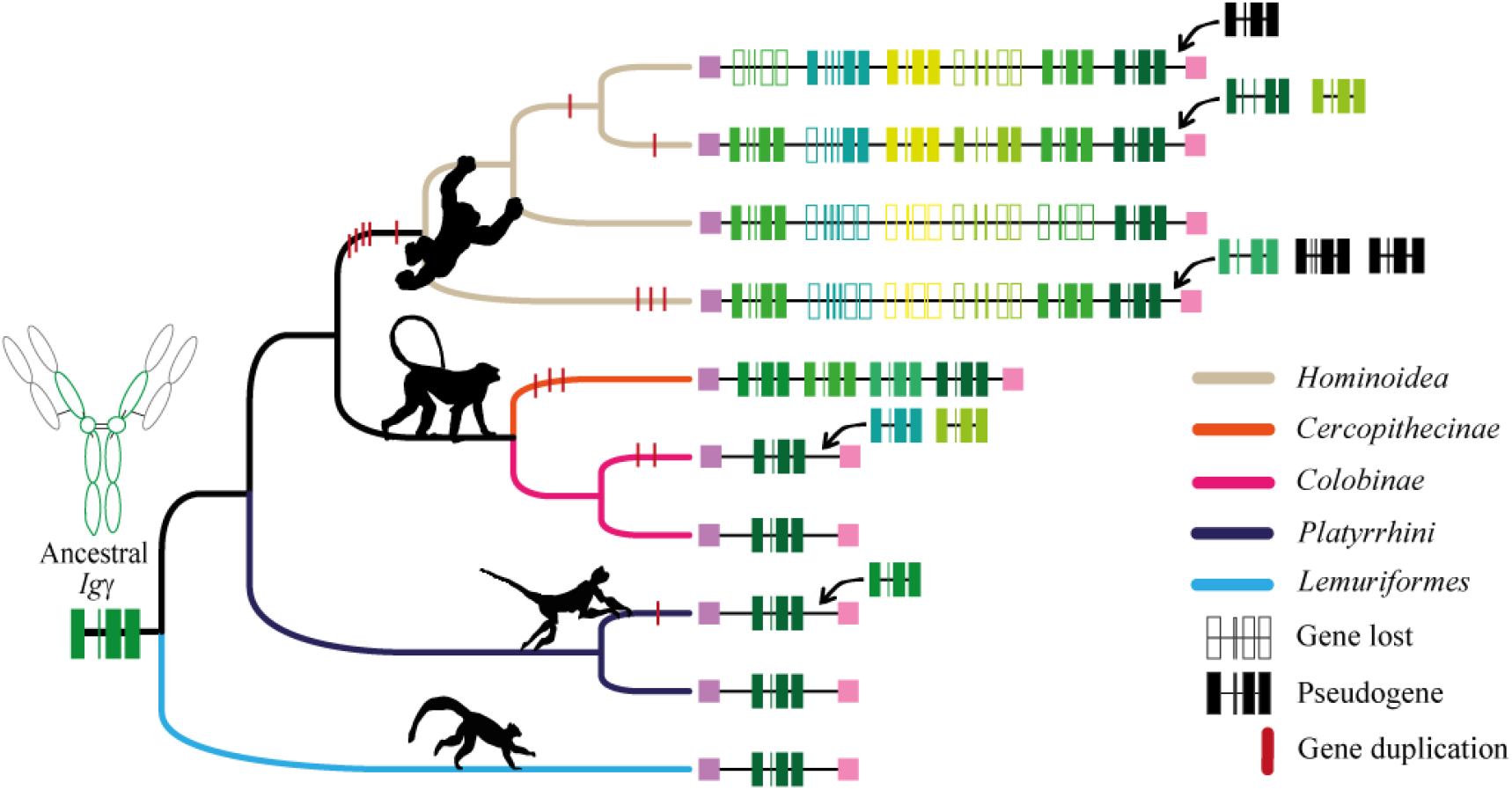

## 1. Introduction

Antibodies or immunoglobulins (Ig) are the primary molecules mediating humoral immunity. These molecules are found in all gnathostomes (jawed vertebrates), evolving to block or neutralize pathogens (bacteria, viruses, and parasites). They are formed by four polypeptide chains, two immunoglobulin heavy (H) chains and two light (L). The N-terminal or Fab (Fragment antigen-binding) region is extremely diverse and responsible for recognizing and interacting with the antigen, while the C-terminal or Fc (fragment crystallizable region, the “antibody tail”) region is responsible for triggering the effector cell responses. In mammals, from the most primitive (egg-laying mammals) to primates, there are five genes: *mu* (*μ*), *delta* (*δ*), *gamma* (*γ*), *epsilon* (*ε*), and *alpha* (*α*), each encoding a different Ig class (IgM, IgD, IgG, IgE, and IgA, respectively) (Senger et al., 2015). They carry out diverse effector functions, such as complement binding, phagocytic cell binding, opsonization, and transport through the placenta epithelium (Schroeder and Cavacini, 2010).

IgG plays an essential role in mammalian adaptive immunity. Unlike other Ig, class G is involved in activating the decisive effector responses that allow antigen/pathogen neutralization or elimination. Moreover, this is the most abundant Ig in the serum, with a half-life of 23 days (Birdsall, 2015; Vidarsson et al., 2014) and it is the only isotype transmitted through the human placenta to the fetus through the placental transport receptor (FcRn). There are four IgG subclasses in humans, each with unique features and functions, although they share more than 90% amino acid sequence similarity (Nimmerjahn and Ravetch, 2005; Vidarsson et al., 2014). For instance, IgG1 and IgG3 can efficiently trigger the classical complement pathway; however, IgG2 and IgG4 do it less efficiently. IgG1 and IgG3 are also the predominant subclasses involved in responding to protein antigens, while IgG2 participates in responding to polysaccharide antigens. Placental transport is also higher for IgG1 and IgG3 but lower for IgG2 and IgG4. IgG3 has a powerful pro-inflammatory effect, given that it is the first IgG to appear after viral infections, followed by IgG1. On the other hand, higher levels of IgG4 have been associated with asymptomatic infections caused by helminths and filariasis (Mishra et al., 2019; Napodano et al., 2020; Vidarsson et al., 2014). Likewise, IgG abundance in serum varies among subclasses; IgG1 represents up to 70% of the total IgG, followed by IgG2 (up to 30%), and IgG3 (∼8%); IgG4 is the least frequent subclass (less than 5%) (Schur, 1987).

These IgG subclasses are encoded by paralogous *γ* genes resulting from tandem duplication events which has been differentially expanded in mammal species (Esteves and Binaghi, 1972; Sun et al., 2012); Platypus, a primitive mammal, shows two *γ* paralogs genes. Cattle displays three while humans, mice, and rats show four copies. At least six distinct *γ* duplicates have been reported in pigs. To date, the horse is the mammal with the largest copy-number (7 genes) (Sun et al., 2012). The number of *γ* genes also appears be different in primates. Macaques have four paralogs (Jacobsen et al., 2011; Ramesh et al., 2017) while in humans there are five (Takahashi et al., 1982), in contrast to new world monkeys where IgG is encoded by a single-copy gene (Brusco et al., 1998; Garzon-Ospina and Buitrago, 2020; Olivieri and Gambon Deza, 2018). Furthermore, in spite macaques and humans have four functional paralogs there does not appear to be a structural and functional correlation between subclasses (Crowley and Ackerman, 2019; Mishra et al., 2019; Nguyen et al., 2014; Ramesh et al., 2017; Warncke et al., 2012). This suggest that *Igγ* multigene family evolutionary history could differ among primates.

Multigene families can evolve by divergent evolution (duplicates diverged gradually acquiring new functions), concerted evolution (duplicates evolve in a concerted manner rather than independently), or by the birth-and-death model (duplicates may be maintained for a long time, whereas others are deleted or become nonfunctional) (Nei and Rooney, 2005). To the best of our knowledge it is not clear how *Igγ* has expanded or contracted in the primate order. With the purpose of assess the evolutionary history of this gene in these mammals, we screened 38 primates genomes available in the GenBank database and performed phylogenetic and molecular evolutionary analyses. This data can help understand the differences among the primates’ immune system and its evolution to cope with infections. Likewise, it could have implications for biomedical trials using these animals.

## 2. Materials and Methods

### 2.1 Datasets and Igγ gene identification

The genome sequences from 38 primate species (Supplementary material 1) available in GenBank (Benson et al., 2015) were analyzed to identify the DNA regions encoding the immunoglobulin heavy constant gamma gene (*Igγ*). The genomic region containing the constant *Igγ* loci is circumscribed by *Igδ* and *Igε* heavy chain genes. Therefore, for those species with a complete ensemble genome, the chromosome region encompassing both *Igδ* and *Igε* genes were analyzed to identify putative *Igγ* loci. For the remaining species, a Blast search was performed to locate DNA contigs containing *Igγ* genes. To determine the exon-intron gene structure, the putative *Igγ* genes were aligned with human IgG CDS subclasses (GenBank accession numbers: AJ294730.1, AJ294731.1, AK097307.1, and AJ294733.1) using the MUSCLE method (Edgar, 2004). Alignments were manually edited in GeneDoc software (K. and H., 1997), identifying the donor and acceptor-splice sites (GT…AG). The coding DNA sequences found were used to deduce amino acid sequences using the GeneRunner software. These sequences were then screened to distinguish immunoglobulin domains through the Pfam webserver (El-Gebali et al., 2019).

### 2.2. The primate Igγ family’s phylogenetic relationships

*Igγ* gene sequences and their deduced-amino-acid-sequences were aligned using the MUSCLE method. The best DNA and protein substitution models were selected using Akaike information criterion through the JModelTest (Posada, 2008) and ProtTest (Darriba et al., 2011) methods, respectively. The GTR+I+G and JTT+G+F models were used to infer phylogenetic DNA and amino acid trees, respectively, using the Bayesian method (BY). A Metropolis-coupled Markov Chain Monte Carlo (MCMC) algorithm was used for this analysis using MrBayes (Huelsenbeck et al., 2001). It was run for 10 million generations. Sump and Sumt commands were used for tabulating posterior probabilities and building a consensus tree. Both analyses were carried out using the CIPRES Science Gateway (Miller MA, 2010; Miller et al., 2015).

Alternatively, we used the multigene family model called DLTRS (Duplications, losses, transfers, rates and sequence evolution) (Sjostrand et al., 2014). This method infers a gene tree by evolving down on a given species tree through duplication, loss, and transfer events according to a birth-death-like process (Sjostrand et al., 2012; Sjostrand et al., 2014). The species tree was inferred with a concatenate fragment of cytochrome C oxidase subunit I and MT-CO2 pseudogene 9. DLTRS was then run using the MCMC algorithm for 10 million generations.

Once the introns were removed, a multiple DNA alignment for *Igγ* was carried out considering the amino acid sequence information using the MUSCLE method in the TranslatorX webserver (Abascal et al., 2010). This alignment was then used to assess natural selection signatures using Datamonkey (Weaver et al., 2018). The Branch-site Unrestricted Statistical Test for Episodic Diversification (BUSTED) method (Murrell et al., 2015) was used to identify whether the *Igγ* gene had experienced positive selection. This gene-wide test assesses positive selection at least in one site on at least one branch. Then, the adaptive Branch-Site Random Effects Likelihood (aBSREL) method (Smith et al., 2015) was performed to identify those *Igγ* branches (lineages) under episodic diversifying selection. In addition, natural selection signatures at individual codon sites were detected by using codon-based tests (FUBAR (Murrell et al., 2013), MEME (Murrell et al., 2012), and SLAC (Kosakovsky Pond and Frost, 2005)). These tests infer the nonsynonymous (d_N_ or K_N_) and synonymous (d_S_ or K_S_) substitution rates for each site basis for a given coding alignment and corresponding phylogeny (Weaver et al., 2018). A sliding window for omega rates (ω = K_N_/K_S_) was draw using the SLAC data to assess the effect of natural selection throughout the *Igγ* gene sequence. Because recombination can bias K_N_/K_S_ estimation (Anisimova et al., 2003; Arenas M, 2014; Arenas and Posada, 2010), the GARD method (Kosakovsky Pond et al., 2006) was performed before running codon-based tests. Finally, the RELAX method (Wertheim et al., 2015) was performed. It allows partitioning a codon-based phylogeny into two subsets of branches to assess whether selective strength was relaxed or intensified in one of these subsets (test branch) relative to the other (reference branch).

### 2.3. Estimation of Igγ paralogs divergence times

The 4-fold degenerate sites from the *Igγ* coding alignment were used to estimate the divergence times among *Igγ* paralogs using the Bayesian MCMC approach available in BEAST software (Drummond and Rambaut, 2007; Suchard et al., 2018). BEAUti (Drummond and Rambaut, 2007; Suchard et al., 2018) was used to generate a BEAST XML file. Divergence times were estimated for individual species using the evolutionary model GTR+G with the relaxed clock option and simulating a Yule process. Four calibration dates (*Cebus albifrons*/*Cebus capucinus*: 1.89 +/- 0.1; *Eulemur macaco*/*Eulemur fulvus*: 7.66 +/- 0.14; *Pithecia pithecia*/*Plecturocebus donacophilus*: 18 +/- 0.6 and *Indri indri*/*Propithecus coquereli*: 29 +/- 6.6) were used based on Timetree web server data (http://www.timetree.org/ (Kumar et al., 2017)). The MCMC length chain was run for 50 million generations. Tracer v1.5 (Rambaut et al., 2018) was used to check when the MCMC chains had reached a stationary distribution by visual inspection of plotted posterior estimates and by checking that ESS (Effective Sample Size) was greater than 1000 for divergence times.

## 3. Results

### 3.1. Igγ gene identification in primate genomes

The sequences containing the immunoglobulin heavy constant gamma locus from 38 primate species were screened to describe the number of putative genes encoding IgG in these species. *Strepsirrhini* and *Haplorhini* suborders displayed differences in the *Igγ* copy number (Fig. 1). A single *Igγ* gene was found in all *Lemuriformes* (*Strepsirrhini*). As mentioned previously, in most *Platyrrhini* species, IgG seems to be encoded by a single-copy gene. *Alpa* was the only species in this parvorder containing two *Igγ* copies (paralogs) (Garzon-Ospina and Buitrago, 2020). On the other hand, the *Catarrhini* species had a high copy-number diversity. Some *Cercopithecoidea* species showed just one *Igγ* gene; other species had three or four duplicates; this feature also was detected in *Hominoidea. Gogo* was the species with the lowest *Igγ* copy-number, displaying two paralogues, followed by *Hosa* with five, *Poab* with six, and *Patr* with eight duplicates. All *Igγ* copies were named in ascending order according to their position in the contigs from left to right from *Igδ* gene. The *Hosa Igγ* subclasses were numbered as is typically done, *Igγ3, Igγ1, IgγΨ, Igγ2*, and *Igγ4*, respectively.

**Figure 1.**
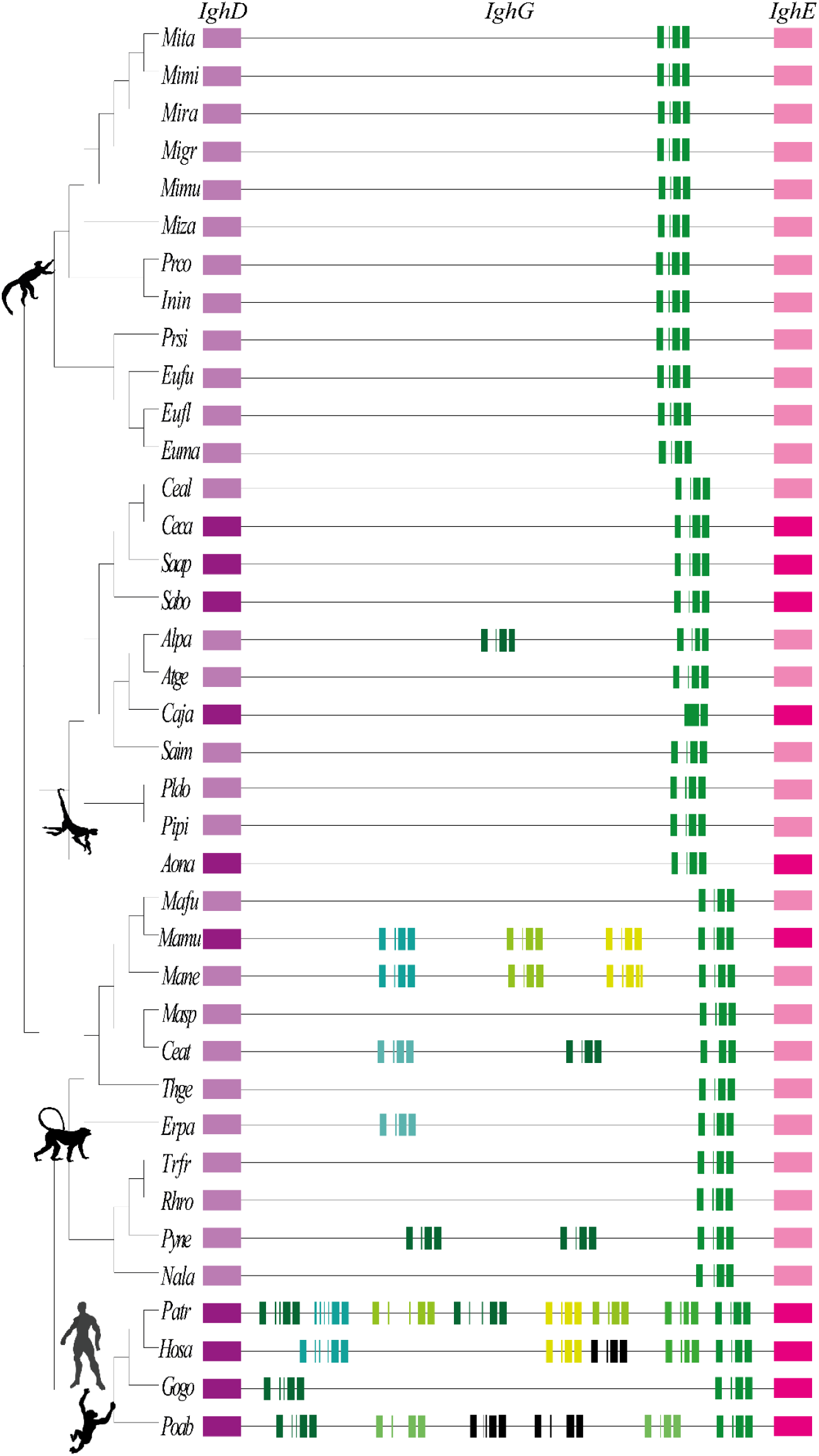
Schematic representation of the chromosomal *Igγ* loci in 38 primate genomes. The *δ* and *ε* genes flanking the *γ* chromosome locus are represented by purple and fuchsia boxes, respectively. Light purple and fuchsia boxes represent the expected *δ* and *ε* genes since these genomes are not fully sequenced or properly ensembled. *γ* genes are depicted in shades of green while pseudogenes are displayed in black. Structure, size and number of exon-intron for each *Igγ* is represented to scale. However, the length of intergenic regions is not informative.

The *Igγ* exon-intron structure was stable in *Lemuriformes, Platyrrhini*, and *Cercopithecoidea* lineages displaying four exons (Fig. 1 and Supplementary material 2). Although *Caja* had been predicted as single-exon gene (Garzon-Ospina and Buitrago, 2020), a re-analysis suggested that two exons might encode IgG in this species; further cDNA analysis will be required to confirm this. According to the Pfam search, the putative IgG proteins in all these species showed the typical three Ig-domains. In *Hominoidea*, some paralogues have expanded the second exon producing a large hinge encoding region. On the other hand, the *PatrIgγ2* gene seems to have lost the first exon (Fig. 1); therefore, this gene would be encoding a two Ig-domains protein. Finally, *PoabIgγ3* and *PoabIgγ4* had several premature stop codons.

### 3.2. The primate Igγ family’s phylogenetic relationships

Phylogenetic relationships for the *Igγ* family were inferred by Bayesian (BY) and DLTRS methods using both DNA and deduced-amino-acid-sequences (Fig. 2-3 and Supplementary material 3-4). A multiple alignment for 67 *Igγ* gene sequences was used for inferring a BY phylogenetic tree using the GTR+I+G model. This phylogeny showed posterior probabilities higher than > 0.95 for most of the nodes, thus, supporting the branching pattern. Four major clades were observed resembling species relationships. The first *Igγ* clade included sequences from *Strepsirrhini* species. The *Platyrrhini* parvorder formed the second clade, where *Alpa* duplicates were clustered in the same branch. The third clade grouping the genes of the *Cercopithecoidea* superfamily. Two different subclades were observed in this group. The subclade 1 clustered the single gene found for *Colobinae* (*Nala, Trfr, Rhro*, and *Pyne*) and the *Igγ1* from *Cercopithecinae* (*Erpa, Thge, Masp, Ceat, Mane, Mafu*, and *Mamu*). The second subclade was formed by the *Igγ2, Igγ3*, and *Igγ4* identified in *Cercopithecinae* species. However, the *Igγ2* and *Igγ3* genes, from *Pyne* and *Ceat*, showed a species-specific clustering (Fig. 2). The latter mayor clade included paralog genes found in the *Hominoidea* superfamily (Fig. 2). Most of the *Patr* paralogs were clustered together with *Hosa Igγ* genes, whereas *Poab* paralogues were found forming a specie-specific subclade. The same topology was displayed by amino acid sequences BY phylogenetic tree inferred with the JTT+G+F model (Supplementary material 3).

**Figure 2.**
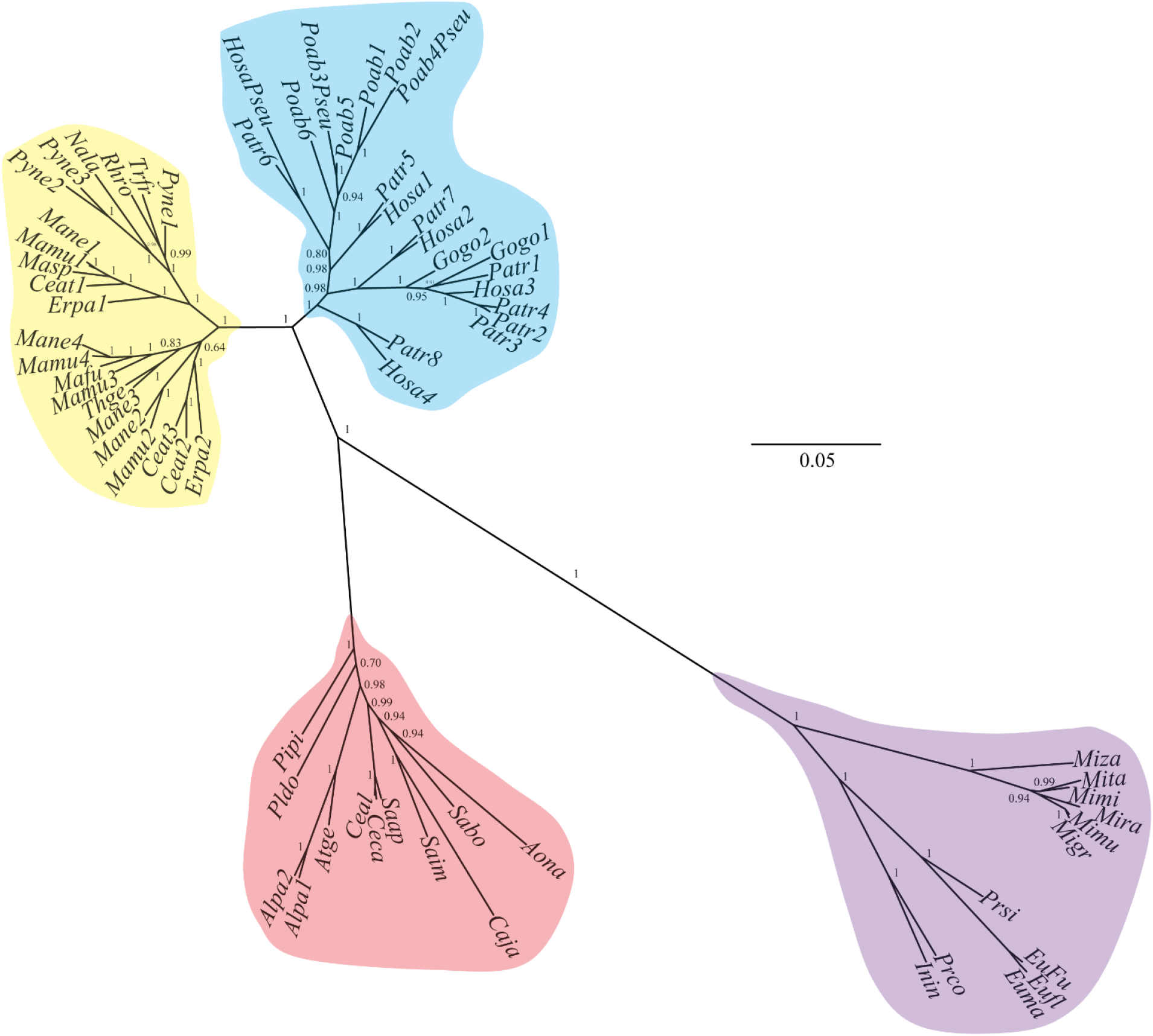
Bayesian (BY) tree inferred for the 67 sequences from the multigene *Igγ* family. Four major clades were identified clustering DNA sequences in agreement with primate phylogenetic relationships. The branching pattern shows posterior probabilities higher than > 0.95 (numbers on branches) supporting the topology. Purple clade cluster the *Strepsirrhini Igγ* gene sequences; red clade put together *Platyrrhini* genes; yellow clade group *Cercopithecoidea* paralogues and the clade clustering *Igγ* duplicates from *Hominoidea* are depicted in blue.

**Figure 3.**
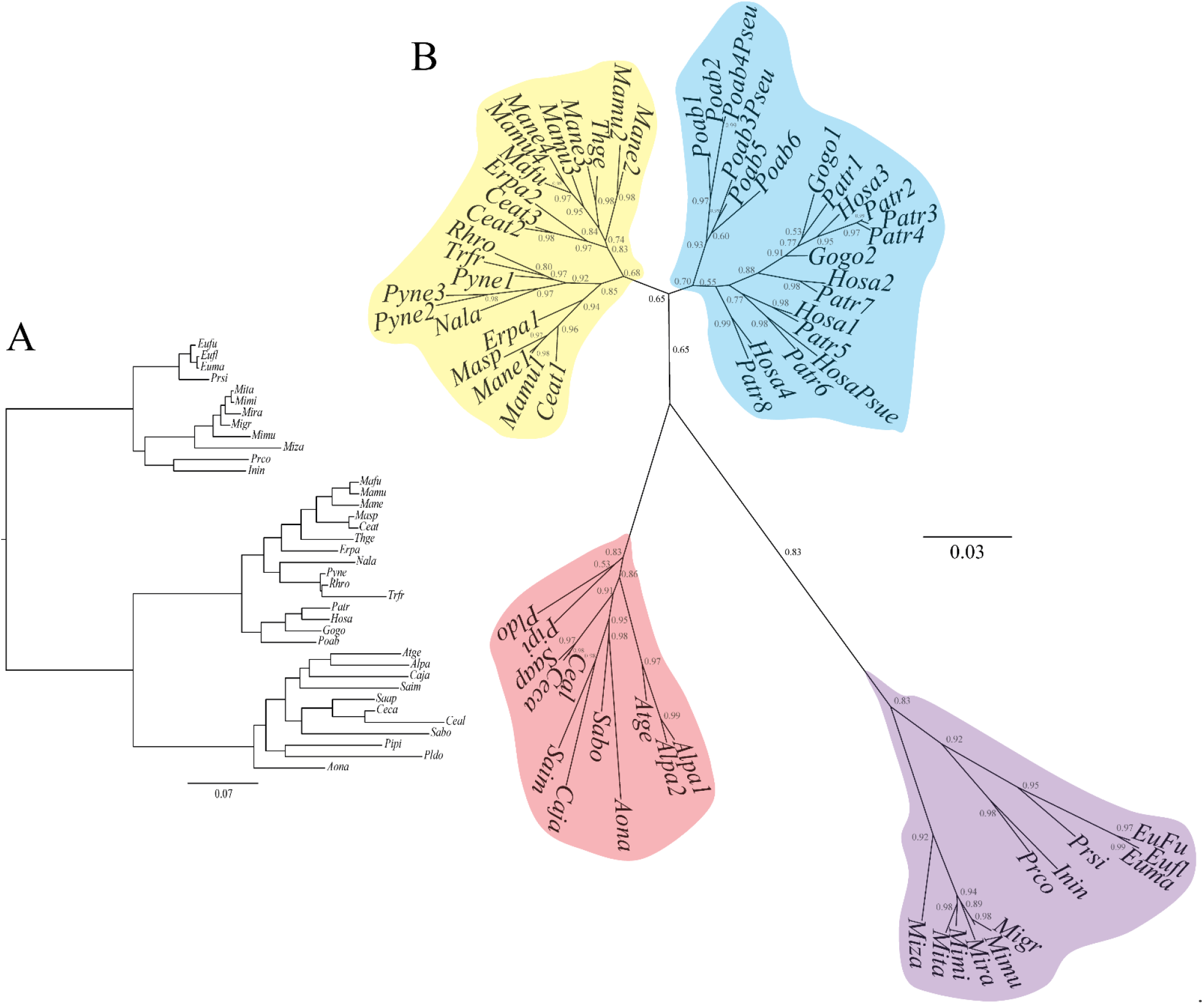
*γ* gene family phylogeny inferred by the DLTRS evolutionary model. Both BY and DLTRS trees showed similar topologies. A. Species tree used for generating the *γ* gene tree. B. *γ* gene tree inferred by evolving down the species tree. Numbers on branches are posterior probability values. Purple clade cluster the *Strepsirrhini Igγ* gene sequences; red clade put together *Platyrrhini* genes; yellow clade group *Cercopithecoidea* paralogues and the clade clustering *Igγ* duplicates from *Hominoidea* are depicted in blue.

In addition, the DLTRS model, which reconciles the gene tree to the species tree (Sjostrand et al., 2012; Sjostrand et al., 2014), was also performed. The trees inferred by DLTRS (for both DNA and deduced-amino-acid-sequences) also had four major clades (Fig. 3 and Supplementary material 4), showing similar branching patterns to the BY topologies. Thus, both methods displayed the same phylogenetic relationships (Fig. 2-3 and Supplementary material 3-4).

### 3.3. Natural selection signatures in Igγ

Several methods were applied to assess how natural selection has occurred in the *Igγ* family. The BUSTED method found evidence of gene-wide episodic diversifying selection in *Igγ* phylogeny (p-value = 0.000 ≤ 0.05). The aBSREL method was performed to identify those *Igγ* branches (lineages) under episodic diversifying selection; 15 branches displayed positive selection signatures (Fig. 4). Thirteen percent of the sites from the branch leading *Haplorrhini* displayed a ω = 8.12. Likewise, the *Platyrrhini* lineage showed episodic selection (ω = 18.1 (7.5% of the sites)). Moreover, several branches showed evidence of episodic diversifying selection within this parvorder.

**Figure 4.**
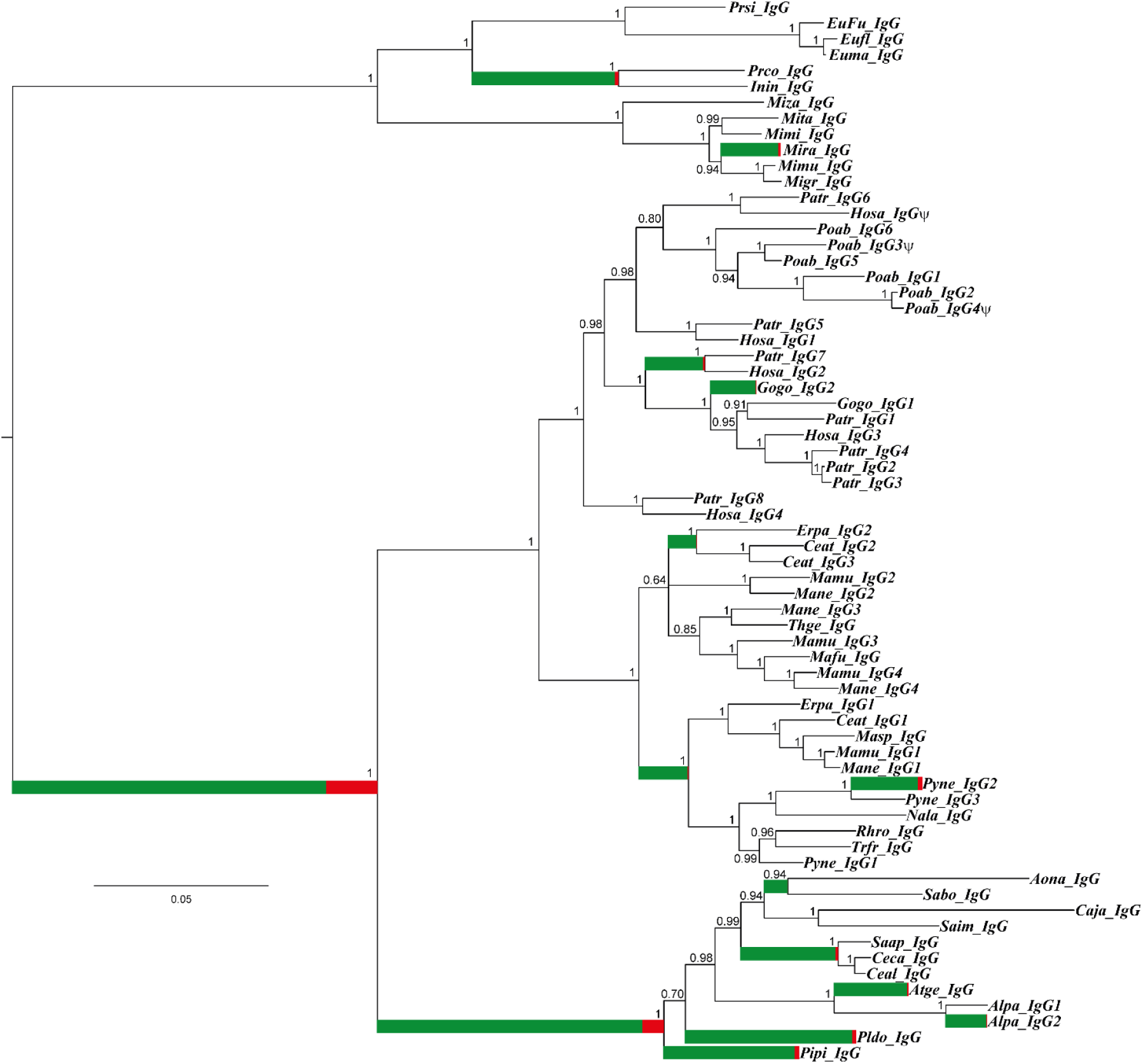
Identification of branches under positive selection. BY tree was analyzed by the aBSREL method. The shade of each color on branches indicates selection strength; red indicates positive selection (ω > 1) and green negative selection (ω < 1). The size of each colored segment represents the percentage of selected sites in the corresponding ω class. Branches have been classified as undergoing episodic diversifying selection, by the p-value corrected for multiple testing, using the Holm–Bonferroni method at p < 0.05.

Codon-based tests identified 77 sites under negative selection (ω < 0) and 44 under positive selection (ω > 0) (Fig. 5 and Supplementary material 5). The negative sites selected were found throughout the *Igγ* gene. Conversely, most of the positive sites selected were located on the CH1 and CH2 domains. Furthermore, the sites homolog to those involved in binding to complement and/or effector cell receptors previously reported for humans displayed a high divergence among species; some of them had signals of positive selection (Fig. 5 and Supplementary material 6).

**Figure 5.**
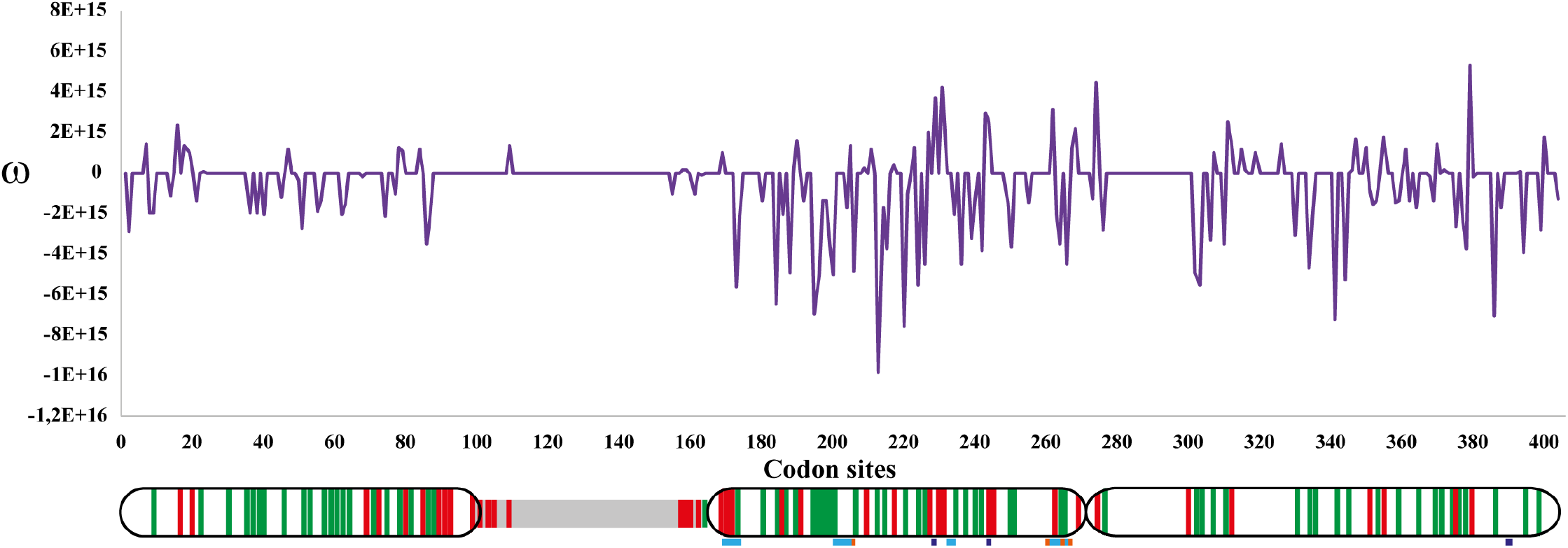
Sliding window for the ω rate. Primate *Igγ* evolutionary rate (ω: K_N_/K_S_) was draw based on SLAC data. A diagram of IgG protein is given below the sliding window. Positive selected sites (red) and negative selected sites (green) are shown by vertical lines within Ig domains (CH1, CH2 and CH3) or hinge region (grey box). Location of sites involved in binding to complement and/or effector cell receptors reported for human are shown as color boxes below protein scheme; IgG-Fc receptors binding sites are displayed in blue, orange box represent the sites involved in complement component 1q binding while neonatal Fc receptor binding sites are showed in dark blue.

Finally, the RELAX method was performed to determine whether selective strength was relaxed or intensified in the branch subsets. Because *Igγ* was not expanded in *Strepsirrhini*, this branch was used as the reference. Both *Platyrrhini* and *Hominoidea* (*Gogo/Hosa/Patr*) showed that intensified selection appears to modulate *Igγ* paralog evolution. In contrast, *Poab* displayed a statistically significant k value lower than 1, indicating a relaxed selection (Supplementary material 7), likely, because two of the six *Poab Igγ* were pseudogenes.

### 3.4. Estimation of Igγ paralogs divergence times

Traditionally, four-fold degenerate sites have been considered to follow the neutral expectation because they are virtually free of selective constraint. Therefore, these sites were used for paralogues divergence time estimation (Table 1 and 2). The paralogues divergence times were consistent with the split times for mayor groups. For instance, according to data on *Hosa* and *Patr* the *Igγ* paralogues divergence took place around 18.45 – 20.29 MYA, which is posterior to the *Cercopithecoidea* and *Hominoidea* split (27.9 – 31.3 MYA (Kumar et al., 2017)) but before *Hominoidea* divergence (14.7 – 16.7 MYA (Kumar et al., 2017)). Likewise, *Mamu* and *Mane Igγs* divergence occurred 21.3 – 22.2 MYA, which is the time estimated for the *Cercopithecoidea* split (16.4 – 22.2 MYA (Kumar et al., 2017)). It is worth noting that some species showed low divergence times between some *Igγ* paralogues. Besides *Alpa*, which was the only *Platyrrhini* with a paralog gene (duplication could have taken place around 2.9 MYA), *Ceat, Pyne, Patr*, and *Poab* also had paralogues that diverged relatively recently (2 – 6 MYA) (Table 1 and 2).

**Table 1.** Paralogues divergence times in *Hominidae*.

| *Hosa* |  | *Patr* |  | *Gogo* |  | *Poab* |  |
| --- | --- | --- | --- | --- | --- | --- | --- |
| *Igys* | Divergence times | *Igys* | Divergence times | *Igys* | Divergence times | *Igys* | Divergence times |
|  |  | 4/3 | 2.05 (0.01) |  |  | 2/ Ψ4 | 1.67 (0.01) |
| Ψ/1** | 14.48 (0.05) | 6/5** | 9.80 (0.03) |  |  | 1/2-Ψ4 | 5.79 (0.02) |
| Ψ/3-1 | 16.36 (0.05) | 4-3-2/1 | 11.53 (0.05) |  |  | 5/ Ψ3 | 5.89 (0.02) |
| 4-2/3-1-Ψ** | 17.02 (0.05) | 8-7/6-5** | 18.96 (0.05) |  |  |  |  |
|  |  | 8/1* | 19.18 (0.06) | 2-1* | 7.97 (0.02) | 6/1* | 11.20 |
| 4-2/3-1§ | 17.26 (0.05) | 8-7/4-3-2§ | 19.21 (0.06) |  |  |  |  |
| 4/2*‡ | 12.87 (0.05) | 8/7*‡ | 19.24 (0.06) |  |  | 6/5‡ | 11.59 |
| *Hosa Igγ* | 18.45 (0.05) | *Patr Igγ* | 20.29 (0.06) |  |  | *Poab Igγ* | 12.54 (0.03) |
|  | *Hosa/Poab*<sup>a</sup> |  | 14.7 - 16.7 |  |  |  |  |
|  | *Hosa/Gogo*<sup>a</sup> |  | 8.38 - 9.58 |  |  |  |  |
|  | *Hosa/Patr*<sup>a</sup> |  | 5.10 - 11.8 |  |  |  |  |

\* and \*\*: Putative orthologs
†, ‡, §, †‡ Putative pseudoorthologs, they had similar location at the genome as well as exon-intron structures.
<sup>a</sup>: Divergence times based on timetree web server data (http://www.timetree.org/ (Kumar et al., 2017)).

**Table 2.**
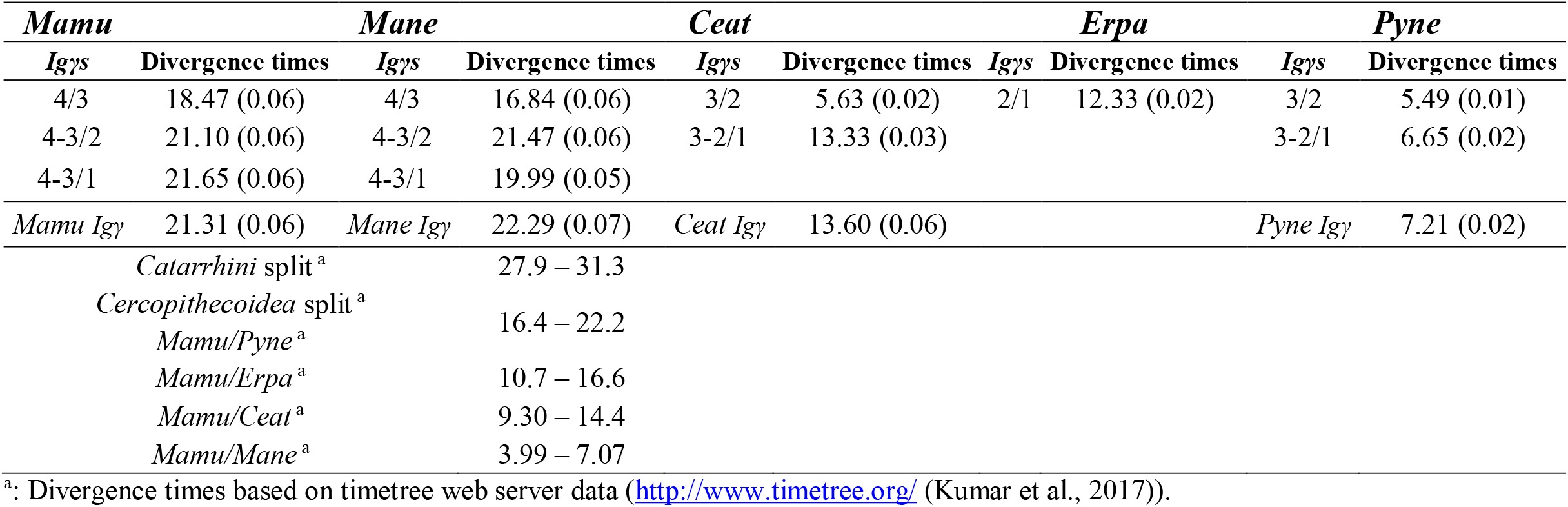
Paralogues divergence times in *Cercopithecoidea*.

## 4. Discussion

Gene duplication is the raw material for evolution (Lynch and Conery, 2000; Stephens, 1951; Zhang, 2003). After duplication, one of the genes (a paralog) can undergo the relaxation of selective strength, allowing the accumulation of mutations that can drive gene inactivation (pseudogenization) or result in acquiring a new function (neofunctionalization) conferring an adaptive advantage (Kondrashov and Kondrashov, 2006; Kondrashov et al., 2002; Krakauer and Nowak, 1999; Zhang, 2003). A multigene family will be formed if duplication is repeated several times in the same DNA region.

Multigene families can evolve by divergent evolution, concerted evolution, or by the birth-and-death model (Nei and Rooney, 2005). The latter appears be common in vertebrates adaptive immune system (Hughes and Nei, 1990; Nei et al., 1997; Ota and Nei, 1994; Su and Nei, 2001). In fact, the main molecules mediating humoral immune responses have emerged by gene duplication. Some of them evolve by a birth-and-death process (Nei et al., 1997; Ota and Nei, 1994; Senger et al., 2015). It has previously been shown that immunoglobulin heavy constant gamma gene (*Igγ*) number varies greatly among *Platyrrhines* and *Catarrhini* (Brusco et al., 1998; Garzon-Ospina and Buitrago, 2020; Olivieri and Gambon Deza, 2018), suggesting that IgG evolution was different among primates.

The *Igγ* phylogenetic trees inferred showed a branching pattern that resembles the species’ phylogeny, suggesting that this gene has evolved in a linage and/or species-specific manner. The 12 *Lemuriformes* and 10 *Platyrrhines* species displayed a single *Igγ* gene in their genomes, suggesting that this region has remained stable for more than 20 million years in these primates. *Alpa* was the only one having a paralog gene that appears to have emerged recently.

On the other hand, the evolutionary history of this gene was highly complex in *Catarrhini*. Previous studies have shown that the IgG subclasses effector function appears to be species-specific among humans and macaques (Crowley and Ackerman, 2019; Nguyen et al., 2014; Ramesh et al., 2017; Warncke et al., 2012). For instance, the interaction of IgG2 and IgG4 to FcR receptors is more effective in cynomolgus monkey than in humans (Warncke et al., 2012). However, this would not necessarily be explained by changes in the critical amino acids of the IgG2 and IgG4 subclasses (Warncke et al., 2012), rather because these duplications are lineage-specific and therefore not orthologous. In addition to these lineage-specific duplications, species-specific duplications have also occurred within the *Catarrhini*. In *Cercopithecoidea*, the *Igγ* number varies. Some species had a single-copy gene, while others had three or four paralogs. This behavior might be because the DNA region containing the immunoglobulin *γ* locus has not been fully sequenced yet and thus, some duplicates could be not found in this study. For instance, *Mafu, Mamu*, and *Mane* are closely related species with a split around 3.99 – 7.07 million years ago (Kumar et al., 2017). However, only one gene was found in *Mafu*, whereas four different genes were observed in *Mamu* and *Mane* genomes.

Although not all genes were found in *Mafu*, the data obtained for others *Cercopithecoidea* suggest an *Igγ* differential expansion. *Cercopithecinae* (*Erpa, Thge, Masp, Ceat, Mane, Mafu*, and *Mamu*) and *Colobinae* (*Nala, Trfr, Rhro*, and *Pyne*) are two subfamilies of this group. While *Mamu* and *Mane*, belonging to *Cercopithecinae*, had four genes, *Nala, Trfr*, and *Rhro*, from the *Colobinae*, displayed a single one. It has been previously scanned some primate genomes in order to identify exon sequences encoding immunoglobulin CH domains (Olivieri and Gambon Deza, 2018). The authors found that in *Nala* and *Rhro* genomes, there was only one *Igγ* gene, which is consistent with our findings. Thus, it can be suggested that *Igγ* is a single-copy gene in these species. Therefore, *Igγ* duplication took place after *Cercopithecinae* and *Colobinae* split. On the other hand, *Ceat* and *Pyne* had three genes; *Igγ2* and *Igγ3*, within these species, were more related to each other than to genes from other species. Because they appear to have diverged relatively recently (2-6 MYA), it can be suggested that *Ceat* and *Pyne Igγ* paralogs have emerged by the species-specific duplication process (they are in-paralogs).

In *Hominidae*, species-specific duplications and gene loss were more noticeable. *Gogo* was the species with the lowest *Igγ* gene number with two duplicates, while *Patr* showed eight paralogs. Most of them seem to have emerged before the main ape groups’ divergence (out-paralogs). However, few of them had orthologs in the ape species assessed here. Hence, several duplicates could have been lost in some *Hominidae*-species evolutionary history. For instance, four of the six *Poab Igγ* genes appear not to have orthologs in other *Hominidae* species because they form a monophyletic clade. This could be due to concerted evolution (a process that homogenizes gene duplicates) or species-specific duplication that arose after *Poab* diverged from other apes. Divergence times support the latter scenario. *PoabIgγ*5 and *PoabIgγ*3 split 5.8 MYA while *PoabIgγ*2 and *PoabIgγ*4 diverged 1.6 MYA. After gene duplication, *PoabIgγ*3 and *PoabIgγ*4 became pseudogenes (several premature stop codons were found), ruling out concerted evolution. Meanwhile, *PoabIgγ6* and *PoabIgγ1* could be *Hominidae* gene orthologs (or even pseudoorthologs) to *PatrIgγ*8/*HosaIgγ*4 and *PatrIgγ1*/*GogoIgγ1*, respectively, due to their similarities in exon-intron structures. The remaining ancestral genes (*PatrIgγ2*/*HosaIgγ*3, *PatrIgγ*5-6/*HosaIgγ1-Ψ* and *PatrIgγ3-4*) were lost in *Poab*.

*Gogo* also appears to have lost most *Igγ* orthologs because only two genes were located in its genome. Previous reports could support this hypothesis. (Olivieri and Gambon Deza, 2018) found 11 IgG encoding*-*exons in *Igh* locus. The first one was located just after the *Igδ* gene; the next three exons formed the first functional gene. Another exon was found next to the *Igε* paralog, while the last six exons (belonging to two different genes) were upstream of the *Igε*-*Igα* genes. However, one gene seems to have been inserted inside another, leaving only one of them as a functional gene (Olivieri and Gambon Deza, 2018). Therefore, most of the genes that emerged early in *Hominidae* evolution were lost or inactivated in this species.

Conversely, several *Igγ* orthologs were shared between *Hosa* and *Patr*. However, in *Hosa*, some ancestral genes were also lost. In this species, no gene showed the same exon-intron structure as *PatrIgγ1*/Gogo*Igγ1*/*PoabIgγ1*. Therefore, this gene seems to have been maintained in most ape species but lost in *Hosa*. Likewise, the *PatrIgγ4-3* lineage was also lost in *Hosa*.

Regarding *Patr*, the *PatrIgγ2* gene (which could have given origin to *PatrIgγ4-3* lineage) was put together with the *HosaIgγ3* in the phylogenetic trees. Both of them displayed a similar exon-intron structure, suggesting that they are orthologues. However, *PatrIgγ2* lost the first 842pb. It is unclear whether this is a pseudogene in this species, or it encode for an IgG with just two immunoglobulin domains. Finally, *PatrIgγ4* and *PatrIgγ3* were species-specific duplicates, splitting around 2 MYA.

### 4.1. Natural selection could have favored IgG paralogue neofunctionalization

The outcomes before gene duplication might be subfunctionalization or neofunctionalization. The latter could be facilitated by functional constraints relaxation, followed by positive selection (Lynch and Conery, 2000; Wertheim et al., 2015). However, the number of sites involved in a new function is usually small (Gu, 2003; Newcomb et al., 1997; Perutz, 1984; Zhang et al., 1998) and, therefore, difficult to trace. Moreover, the positive selection acting at these sites could be transient or episodic, making its detection more problematic. However, if a gene/protein has experienced positive selection for a new or modified function, it could be expected that the lineage subject to positive selection shows a higher evolution rate than other phylogenetic lineages (Fay and Wu, 2003; Messier and Stewart, 1997). Thus, several tests have been developed to assess whether positive selection has occurred on a proportion of lineages (Smith et al., 2015). Furthermore, some codon-based methods are useful to identify sites under episodic selection signatures (Murrell et al., 2012).

Here, several tests displayed that the *Igγ* family has experienced positive selection. Several internal branches displayed signatures of positive selection, which could indicate ancestral diversification in the *Igγ* family. Moreover, several positive selected sites were found by codon-based methods, most of them located in the CH2 domain. Critical amino acid residues, binding to complement and effector cell receptors (Fc receptors), have been previously determined in *Hosa*. They have been located at the CH2 and CH3 domains (Idusogie et al., 2001; Vidarsson et al., 2014; Woof and Burton, 2004). Several of these binding sites were not conserved in the *Igγ* primate alignment. Furthermore, some of them showed signatures of episodic positive selection. Diversity and polymorphism (allotypes) have proven to reduce subclass binding or affinity in *Hosa* IgG, as well as placental transport and protein half-life (Vidarsson et al., 2014). Thus, a change in amino acids, favored by positive selection, could have provoked a functional divergence in the *Igγ* paralogs. IgG respond to different antigen types in humans, having a marked skewing towards one of the four IgG subclasses. Furthermore, these subclasses have different affinities to Fc-receptor or complement binding; thus, they can trigger different effector mechanisms (Irani et al., 2015; Jacobsen et al., 2011; Vidarsson et al., 2014). Accordingly, an *Igγ* expansion, followed by positive selection, could have occasioned a high specialized IgG-response in *Catarrhines*.

Likewise, unequal crossover is an important mechanism for gene expansion (or contraction) (Zhang, 2003). Moreover, selective pressures for an expanded (or contracted) number of genes would be favorably selected in a population (Hood et al., 1980). Consequently, a gene might have an optimal number of copies per genome, depending on the environmental conditions (Garzon-Ospina et al., 2010; Kondrashov and Kondrashov, 2006). Thus, the *Igγ* gene family in primate species could be evolving by a recurrent process of lineage-specific (or species-specific) gene duplication and gene loss (birth-and-death model). According to the data reported here, the *Igγ* family might has expanded (or contracted) until reaching an optimal number of copies in each species’ genome in order to generate novel strategies for pathogens recognition, neutralization or elimination as competitive ELISA assays previously suggest (Asada et al., 2002). Consequently, IgG-based immune responses might be species-specific, and thus protective immune response observed in vaccine trials using primate biomedical models could not be fully extrapolated to humans as lately has been suggested (Crowley and Ackerman, 2019; Garzon-Ospina and Buitrago, 2020; Nguyen et al., 2014; Ramesh et al., 2017; Warncke et al., 2012).

### 4.2. A hypothesis for Igγ family evolution

A model for *Igγ* family evolution in primates is proposed here based on the data obtained (Fig. 6). Primate ancestors had a single *Igγ* gene. About 70.9 – 76.9 million years ago, *Strepsirrhini* and *Haplorrhini* split. In the former, *Igγ* locus has been unchanging without duplication events. Then, 41.0 – 45.7 MYA, the *Platyrrhini*/*Catarrhini* divergence took place. Duplication events had not occurred in most *Platyrrhini* species; however, around 2.9 MYA, an *Igγ* paralog arose in *Alpa*. Meanwhile, after *Cercopithecoidea*’s and *Hominidae*’s divergence, several linage and species-specific duplicates emerged. *Cercopithecinae Igγ1* arose 21.6 MYA, then, *Igγ2* emerged (21.1 MYA), and, finally, *Igγ3* diverge from the ancestral *Igγ4* gene around 18.4 million years ago. It is not clear whether some genes were lost in *Erpa, Thge, Masp*, or *Ceat*. However, in the latter species, a new duplication took place 5.6 MYA. Likewise, two paralogs emerged recently in *Pyne*. Lineage *PyneIgγ3-Igγ2* split from *PyneIgγ1* around 6.6 MYA and *Igγ3* and *Igγ2* diverged 5.4 MYA. In other *Colobinae* species (*Nala, Trfr*, and *Rhro*) IgG was encoded by a single-copy gene.

**Figure 6.**
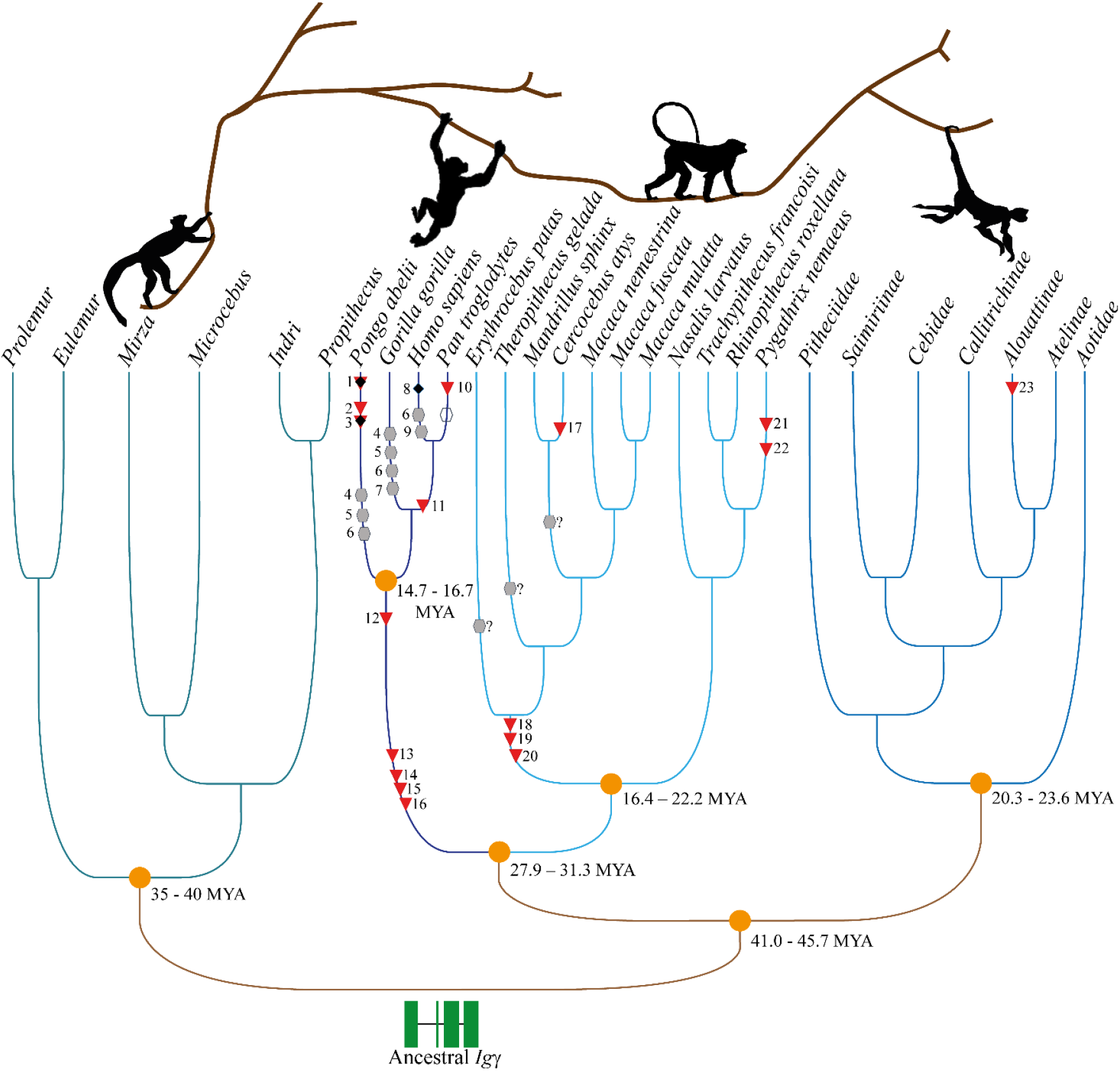
Evolutionary model for the *Igγ* multigene family. Primate ancestor had a single *Igγ* gene. 70.9 - 76.9 million years ago *Strepsirrhini* and *Haplorrhini* split. In *Lemuriformes* as well as *Platyrrhini Igγ* has not undergo duplication. *Alpa* is the only *Platyrrhini* showing a paralog gene. After *Cercopithecoidea* and *Hominidae* divergence (27.9 – 31.3 MYA) several duplicates have emerged. Three duplication events took place in *Cercopithecinae* subfamily whereas in most *Colobinae* species IgG is encoded by a single-copy gene. Recent specie-specific duplication increased *Igγ* copy number in some *Cercopithecoidea* species. *Hominidae* paralogs arose before apes’ divergence however, several of them has been lost or become in pseudogenes. *Hosa* and *Patr* shared several orthologs while *PoabIgγ* duplicates have arose by specie-specific duplication. 1. *PoabIgγ2* and *PoabIgγ4* divergence. 2. *PoabIgγ1* and *PoabIgγ2-4* divergence. 3. *PoabIgγ5* and *PoabIgγ3* divergence 4. Loss of ortholog to *PatrIgγ2*. 5. Loss of ortholog to *PatrIgγ5-6* lineage. 6. Loss of ortholog to *PatrIgγ3-4* lineage. 7. *Gogo* lost the ortholog to *PatrIgγ7/HosaIgγ2*. 8. *PatrIgγ6 ortholog p*seudogenization in *Hosa*. 9. Loss of ortholog to *PatrIgγ*1. 10. *PatrIgγ4* and *PatrIgγ3* divergence. 11. *PatrIgγ6* (*HosaIgγΨ*) and *PatrIgγ5* (*HosaIgγ1*) divergence. 12. Origin of *PatrIgγ3-4* lineage. 13. Origin of *PatrIgγ5-6* (*HosaIgγ1-Ψ*) lineage. 14. Origin of *PatrIgγ1* (*PoabIgγ1, GogoIgγ1*). 15. Origin of *PatrIgγ2* (*HosaIgγ3*). 16. Origin of *PatrIgγ7* (*HosaIgγ2, PoabIgγ5, GogoIgγ*2). 17. *CeatIgγ2* and *CeatIgγ3* divergence. 18. *MamuIgγ3* (*ManeIgγ3*) and *MamuIgγ4* (*ManeIgγ4*) divergence. 19. Origin of *MamuIgγ2* (*ManeIgγ2*). 20. Origin of *MamuIgγ1* (*ManeIgγ1*). 21. *PyneIgγ2* and *PyneIgγ3* divergence. 22. *PyneIgγ2-3* and *PyneIgγ1* divergence. 23. *AlpaIgγ1* and *AlpaIgγ2* divergence. Orange circles represent speciation events; red triangles symbolize duplication events; grey hexagon denote gene lost; unfilled hexagon means loss of the first *PatrIgγ2* exon and black diamond indicate pseudogenization. ?: Loss of genes or absence due to incomplete genomes.

Regarding the *Hominidae* species, the *Igγ* was expanded around 20 MYA. The ancestral gene (*PatrIgγ8/HosaIgγ4/GogoIgγ2/PoabIgγ6*) gave rise to *PatrIgγ7/HosaIgγ2* (19.2 MYA); then, *PatrIgγ2/HosaIgγ3* arose (19.8 MYA). In this gene, the exon two underwent intragenic duplication. Moreover, the *PatrIgγ2* gene lost 842pb towards the 5’-end. Later, the *PatrIgγ5-6/HosaIgγ1-Ψ* lineage emerged around 18.9 MYA, followed by the rise of the *PatrIgγ4-3* lineage about 11.3 million years ago. However, several of these genes were lost before the *Hominidae* split. For instance, in *Poab*, the orthologs to *PatrIgγ2, PatrIgγ5-*6, and *PatrIgγ4-3* were not found at the genome. Nevertheless, in this species, new duplications took place a few million years ago. *PoabIgγ3* diverged from *PoabIgγ5* around 5.9 MYA; then, *PoabIgγ2* and *PoabIgγ4* diverged 1.7 MYA. Both the *PoabIgγ3* and *PoabIgγ4* have become pseudogenes. *Gogo* also lost most of the ancestral duplicates, maintaining just two *Igγ* genes. *Hosa* lost the ortholog to *PatrIgγ1*, as well as the ortholog to *PatrIgγ4-3*. Moreover, *Hosa* orthologue to *PatrIgγ6* was inactivated and is currently a pseudogene. Finally, the last duplication process in this superfamily took place around 2 MYA in *Patr*, giving rise to *PatrIgγ4* and *PatrIgγ3*.

## Abbreviations

*Alpa*: *Alouatta palliate*
*Aona*: *Aotus nancymaae*
*Atge*: *Ateles geoffroyi*
*Caja*: *Callithrix jacchus*
*Ceal*: *Cebus albifrons*
*Ceca*: *Cebus capucinus*
*Ceat*: *Cercocebus atys*
*Erpa*: *Erythrocebus patas*
*Eufl*: *Eulemur flavifrons*
*Eufu*: *Eulemur fulvus*
*Euma*: *Eulemur macaco*
*Gogo*: *Gorilla gorilla*
*Hosa*: *Homo sapiens*
*Inin*: *Indri indri*
*Mafu*: *Macaca fuscata*
*Mamu*: *Macaca mulata*
*Mane*: *Macaca, nemestrina*
*Masp*: *Mandrillus sphinx*
*Migr*: *Microcebus griseorufus*
*Mimi*: *Microcebus mittermeieri*
*Mimu*: *Microcebus murinus*
*Mira*: *Microcebus ravelobensis*
*Mita*: *Microcebus tavaratra*
*Miza*: *Mirza zaza*
*Nala*: *Nasalis larvatus*
*Patr*: *Pan troglodytes*
*Pipi*: *Pithecia pithecia*
*Pldo*: *Plecturocebus donacophilus*
*Poab*: *Pongo abelii*
*Prsi*: *Prolemur simus*
*Prco*: *Propithecus coquereli*
*Pyne*: *Pygathrix nemaeus*
*Rhro*: *Rhinopithecus roxellana*
*Saim*: *Saguinus imperator*
*Sabo*: *Saimiri boliviensis*
*Saap*: *Sapajus apella*
*Thge*: *Theropithecus gelada*
*Trfr*: *Trachypithecus francoisi*

## Data statement

The datasets supporting this article’s results are available in Mendeley Data: Garzón-Ospina, Diego; Buitrago, Sindy P (2020), “Immunoglobulin heavy constant gamma gene evolution is modulated by both the divergent and birth-and-death evolutionary models”, Mendeley Data, V1, doi: 10.17632/r6vh2rfnts.1 (DOI is reserved but not active yet)

## Acknowledgments

The authors wish to thank Gypsy Bonny Español for reviewing the manuscript.

## Supplementary files

**Supplementary material 1. GenBank accession numbers from 38 primate species used here.** *Igγ* genes were searching using genome sequences available in GenBank database.

**Supplementary material 2. Data regarding *Igγ* exon-intron structure.** Exon and intron base pair length for each *Igγ* gene are shown

**Supplementary material 3. Bayesian (BY) tree inferred for the 67 amino acid sequences from the multigene IgG family.** Four major clades were identified clustering amino acid sequences in agreement with primate phylogenetic relationships. The branching pattern shows posterior probabilities higher than > 0.95 (numbers on branches) supporting the topology. Purple clade cluster the *Strepsirrhini* IgG protein sequences; red clade put together *Platyrrhini* genes; yellow clade group *Cercopithecoidea* paralogues and the clade clustering *Igγ* duplicates from *Hominoidea* are depicted in blue.

**Supplementary material 4. IgG protein family phylogeny inferred by the DLTRS evolutionary model.** Both BY and DLTRS trees showed similar topologies using DNA or proteins sequences. A. Species tree used for generating the IgG tree. B. IgG tree inferred by evolving down the species tree. Numbers on branches are posterior probability values. Purple clade cluster the *Strepsirrhini* IgG protein sequences; red clade put together *Platyrrhini* genes; yellow clade group *Cercopithecoidea* paralogues and the clade clustering *Igγ* duplicates from *Hominoidea* are depicted in blue.

**Supplementary material 5. Negative and positive selected sites identified by codon-based methods.** Codon-based methods identified sites under natural selection. Location of these sites within Ig-domains (CH1-CH3) or hinge regions are indicated.

**Supplementary material 6. Alignment of deduced IgG amino acid sequences.** The human IgG critical amino acid residues binding to complement, and effector cell receptors are shown. They are located at the CH2 and CH3 domains. FcγRn (IgG-Fc receptors) binding sites are highlighted in light blue, while yellow sites are those involved in Cq1 (complement component 1q) binding. Yellow asterisks correspond to sites involved in binding to both FcγRn and Cq1. FcRn (neonatal Fc receptor) binding sites are highlighted in dark blue. CH (Ig) domains and hinge region were determined using the EU numbering tool. Positive selected sites within CH2 and CH3 domains are indicated.

**Supplementary material 7. Relaxed or intensified natural selection between reference and tested branches.** Selection intensity (relaxation or intensification) found by the RELAX method is shown for pairwise-lineage comparisons.

## CRediT author statement

**Diego Garzón-Ospina:** Conceptualization, Investigation, Formal analysis, Methodology, Visualization, Writing-Original draft preparation. **Sindy P. Buitrago:** Investigation, Methodology, Visualization, Formal analysis, Writing-Reviewing and Editing

## Highlights

- The evolution of the immunoglobulin gamma has been assessed in primates
- Immunoglobulin gamma has been differentially expanded in primates
- Several linage/specie-specific paralogues were found in primates
- Immunoglobulin gamma in primates evolves by divergent and birth-and-death models

---

**Supplementary material 1.**
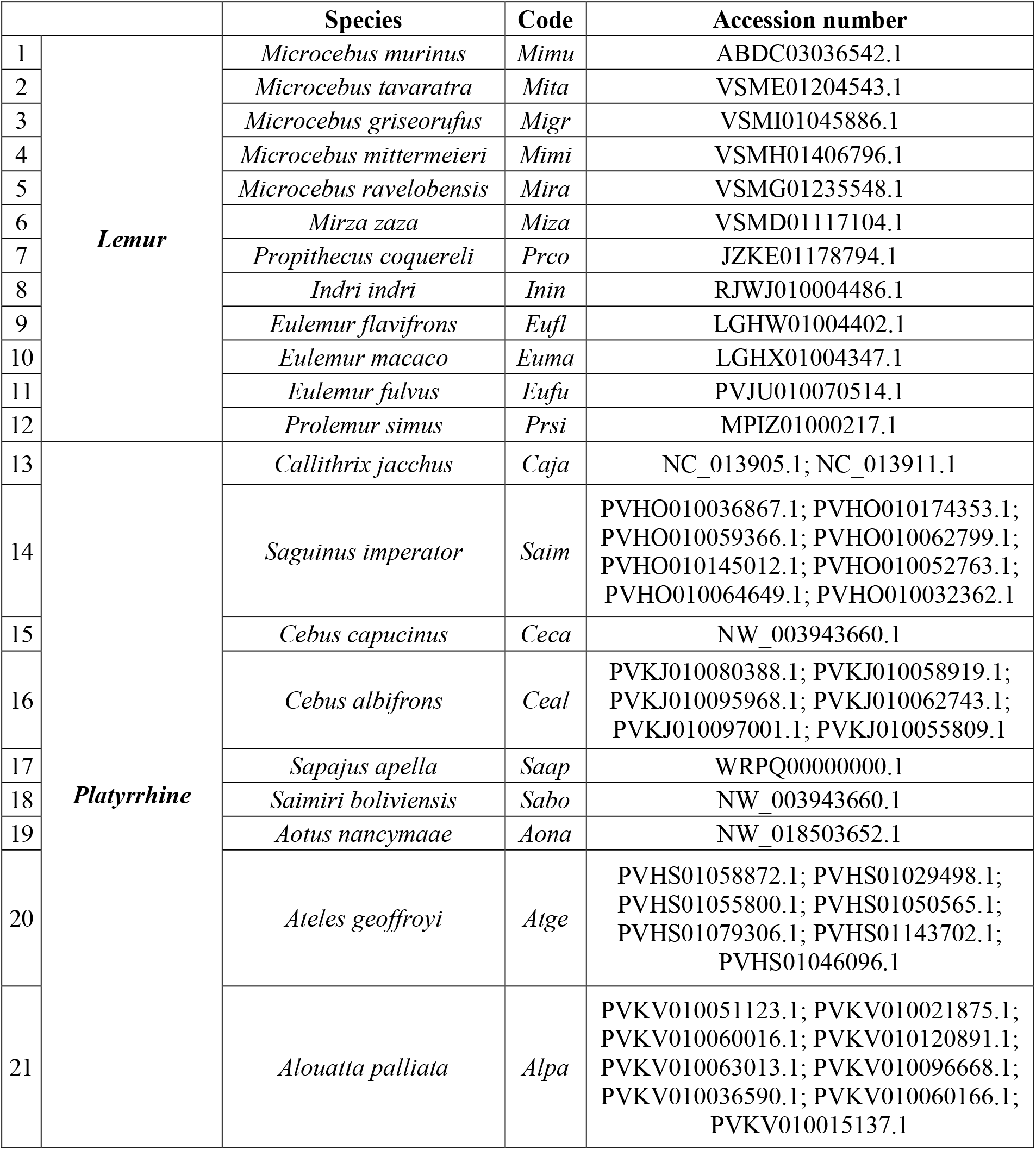
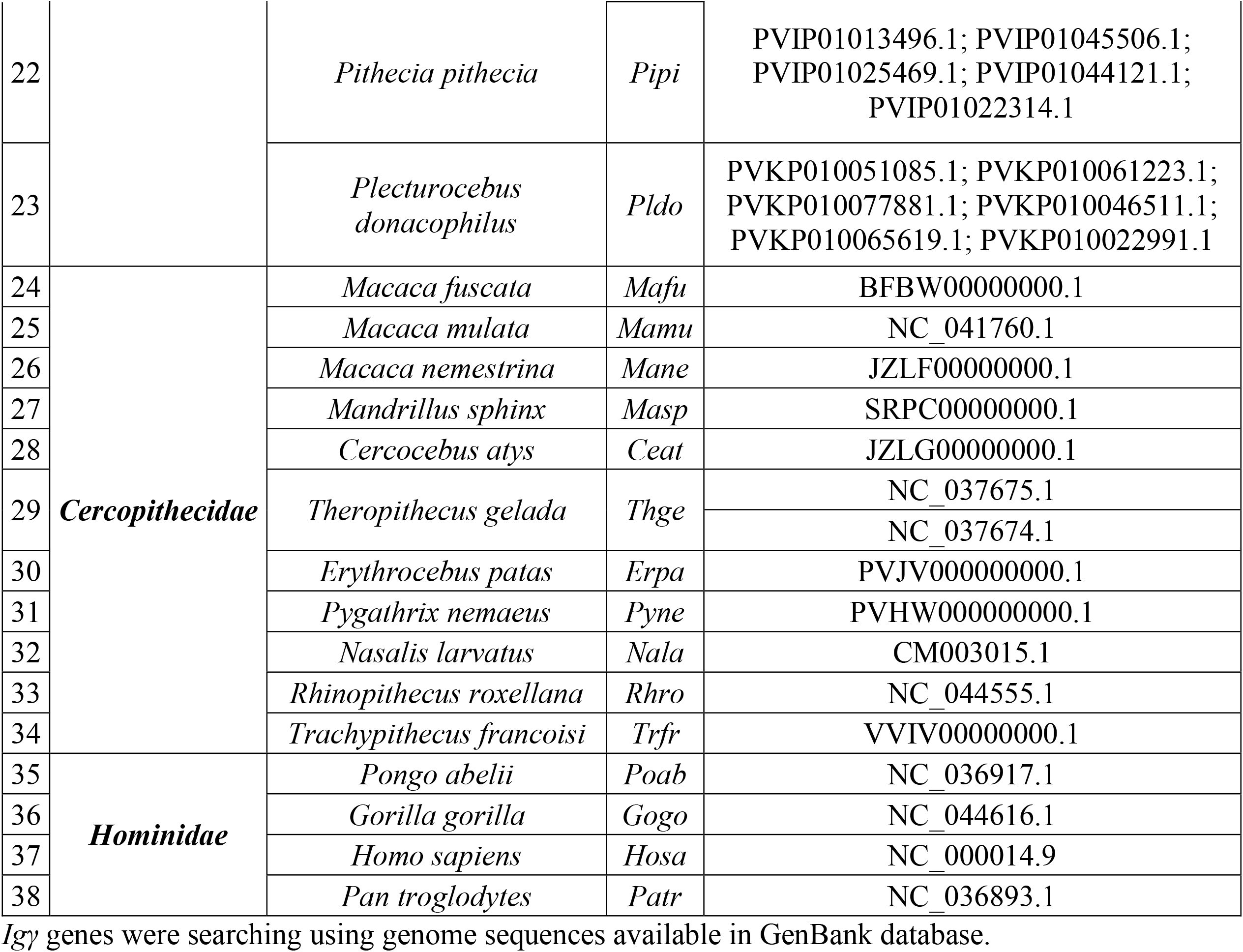
GenBank accession numbers from 38 primate species used here.

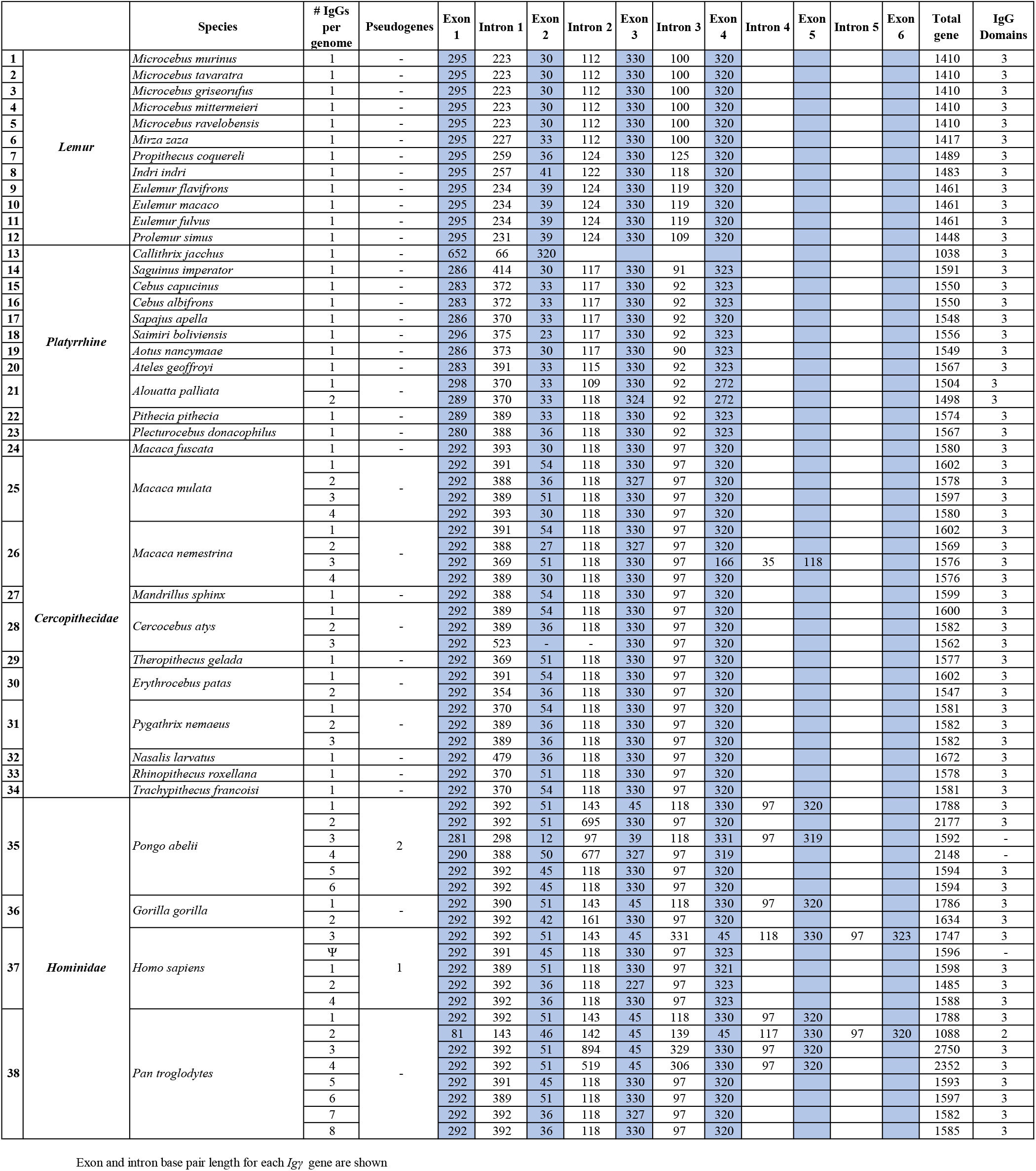

**Supplementary material 3.**
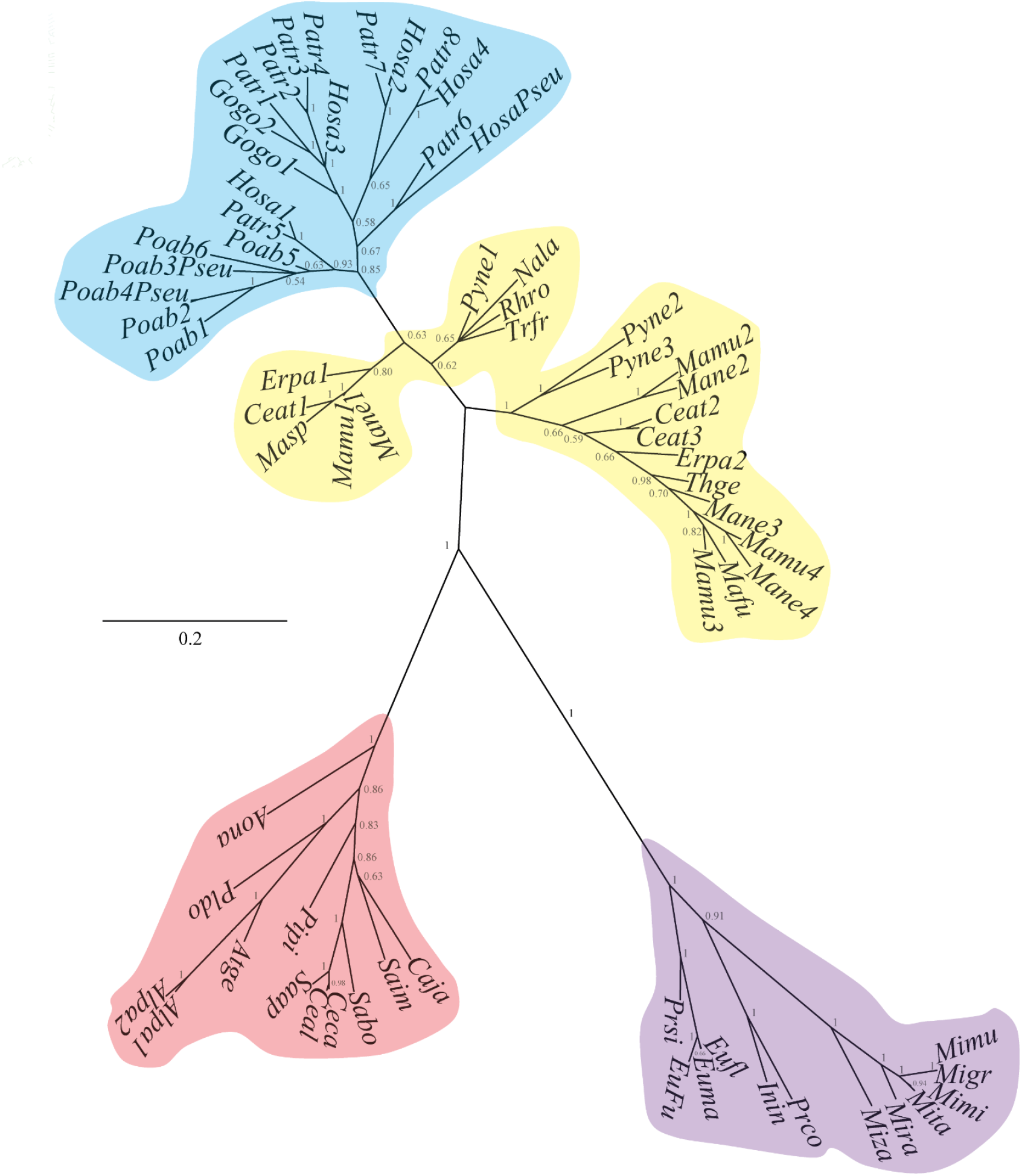
Bayesian (BY) tree inferred for the 67 amino acid sequences from the multigene IgG family

**Supplementary material 4.**
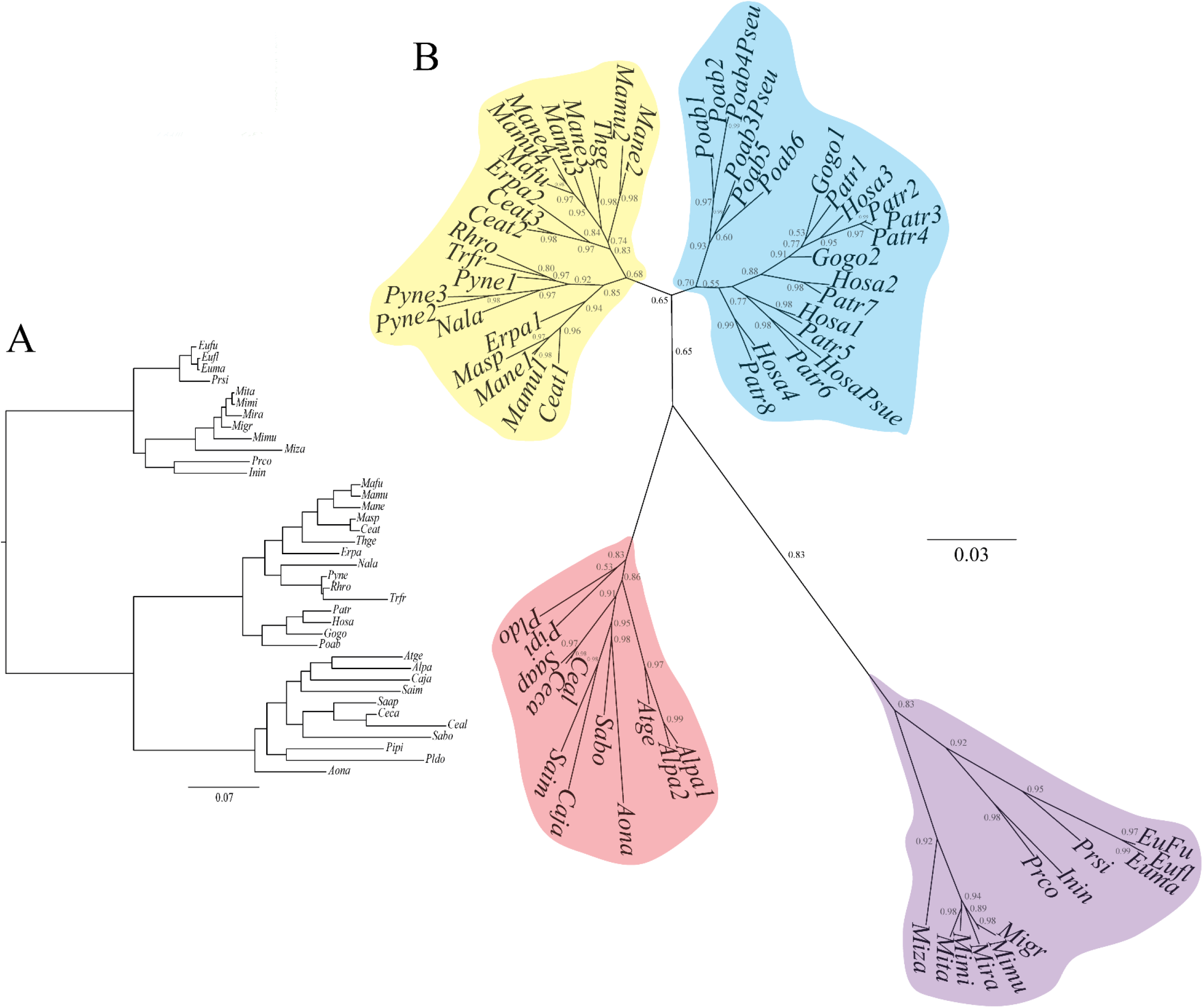
IgG protein family phylogeny inferred by the DLTRS evolutionary model

**Supplementary material 5.**
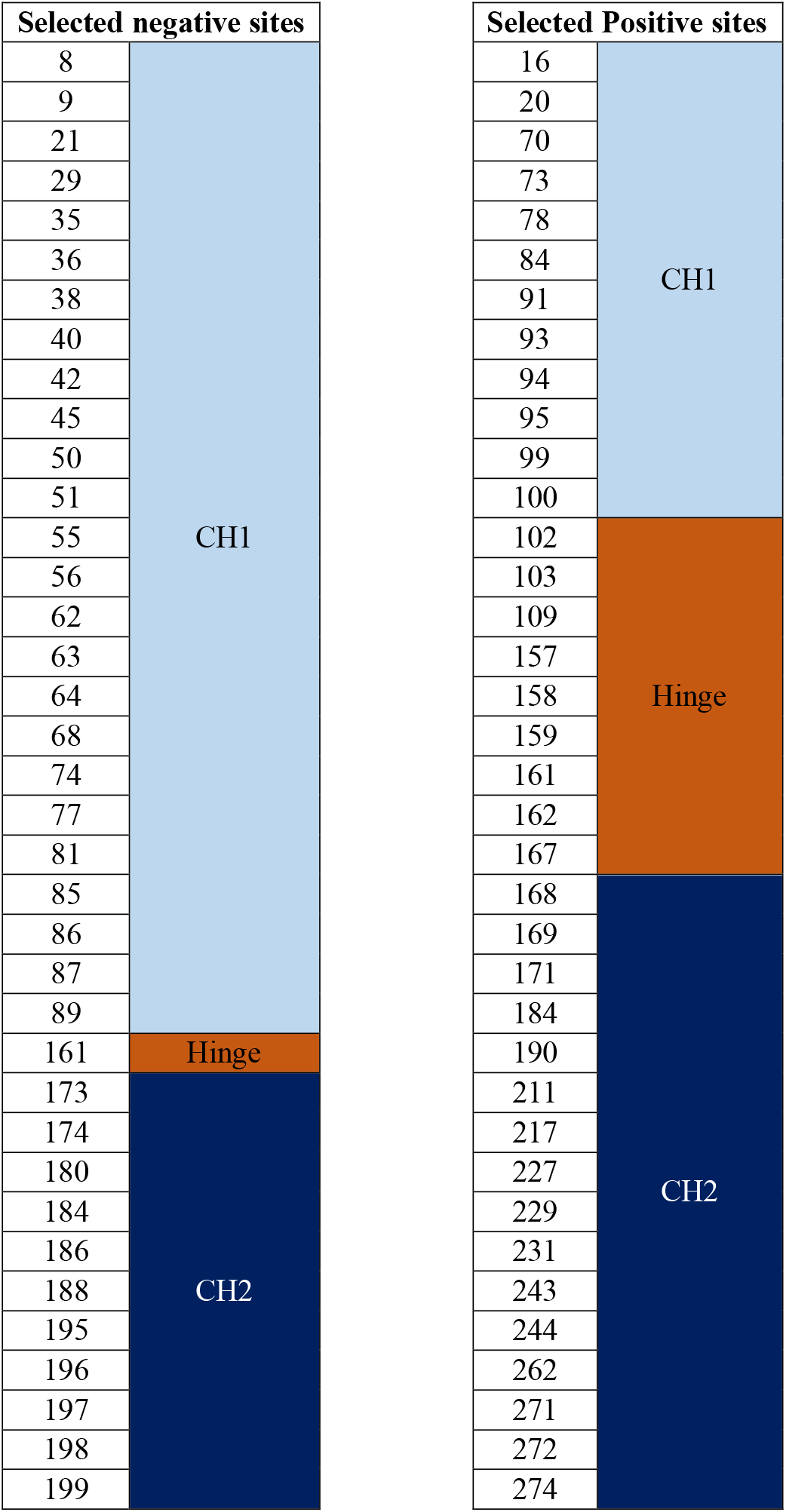
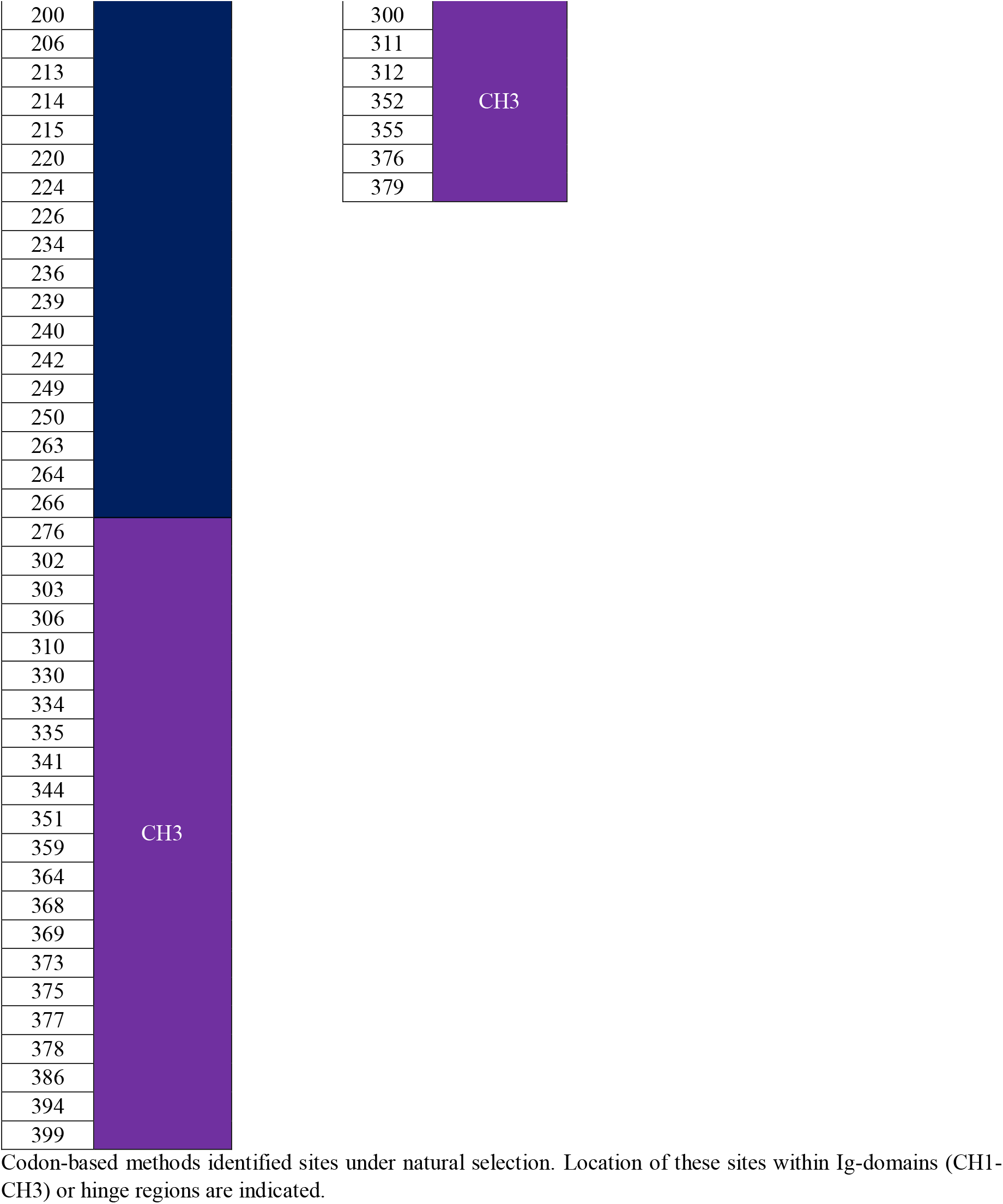
Negative and positive selected sites identified by codon-based methods.

| Selected negative sites |  |
| --- | --- |
| 8 | CH1 |
| 9 |  |
| 21 |  |
| 29 |  |
| 35 |  |
| 36 |  |
| 38 |  |
| 40 |  |
| 42 |  |
| 45 |  |
| 50 |  |
| 51 |  |
| 55 |  |
| 56 |  |
| 62 |  |
| 63 |  |
| 64 |  |
| 68 |  |
| 74 |  |
| 77 |  |
| 81 |  |
| 85 |  |
| 86 |  |
| 87 |  |
| 89 |  |
| 161 | Hinge |
| 173 | CH2 |
| 174 |  |
| 180 |  |
| 184 |  |
| 186 |  |
| 188 |  |
| 195 |  |
| 196 |  |
| 197 |  |
| 198 |  |
| 199 |  |

| Selected Positive sites |  |
| --- | --- |
| 16 | CH1 |
| 20 |  |
| 70 |  |
| 73 |  |
| 78 |  |
| 84 |  |
| 91 |  |
| 93 |  |
| 94 |  |
| 95 |  |
| 99 |  |
| 100 |  |
| 102 | Hinge |
| 103 |  |
| 109 |  |
| 157 |  |
| 158 |  |
| 159 |  |
| 161 |  |
| 162 |  |
| 167 |  |
| 168 | CH2 |
| 169 |  |
| 171 |  |
| 184 |  |
| 190 |  |
| 211 |  |
| 217 |  |
| 227 |  |
| 229 |  |
| 231 |  |
| 243 |  |
| 244 |  |
| 262 |  |
| 271 |  |
| 272 |  |
| 274 |  |

|  |  |  |  |
| --- | --- | --- | --- |
| 200 |  | 300 | CH3 |
| 206 |  | 311 |  |
| 213 |  | 312 |  |
| 214 |  | 352 |  |
| 215 |  | 355 |  |
| 220 |  | 376 |  |
| 224 |  | 379 |  |
| 226 |  |  |  |
| 234 |  |  |  |
| 236 |  |  |  |
| 239 |  |  |  |
| 240 |  |  |  |
| 242 |  |  |  |
| 249 |  |  |  |
| 250 |  |  |  |
| 263 |  |  |  |
| 264 |  |  |  |
| 266 |  |  |  |
| 276 | CH3 |  |  |
| 302 |  |  |  |
| 303 |  |  |  |
| 306 |  |  |  |
| 310 |  |  |  |
| 330 |  |  |  |
| 334 |  |  |  |
| 335 |  |  |  |
| 341 |  |  |  |
| 344 |  |  |  |
| 351 |  |  |  |
| 359 |  |  |  |
| 364 |  |  |  |
| 368 |  |  |  |
| 369 |  |  |  |
| 373 |  |  |  |
| 375 |  |  |  |
| 377 |  |  |  |
| 378 |  |  |  |
| 386 |  |  |  |
| 394 |  |  |  |
| 399 |  |  |  |

Codon-based methods identified sites under natural selection. Location of these sites within Ig-domains (CH1-CH3) or hinge regions are indicated.

**Supplementary material 6.**
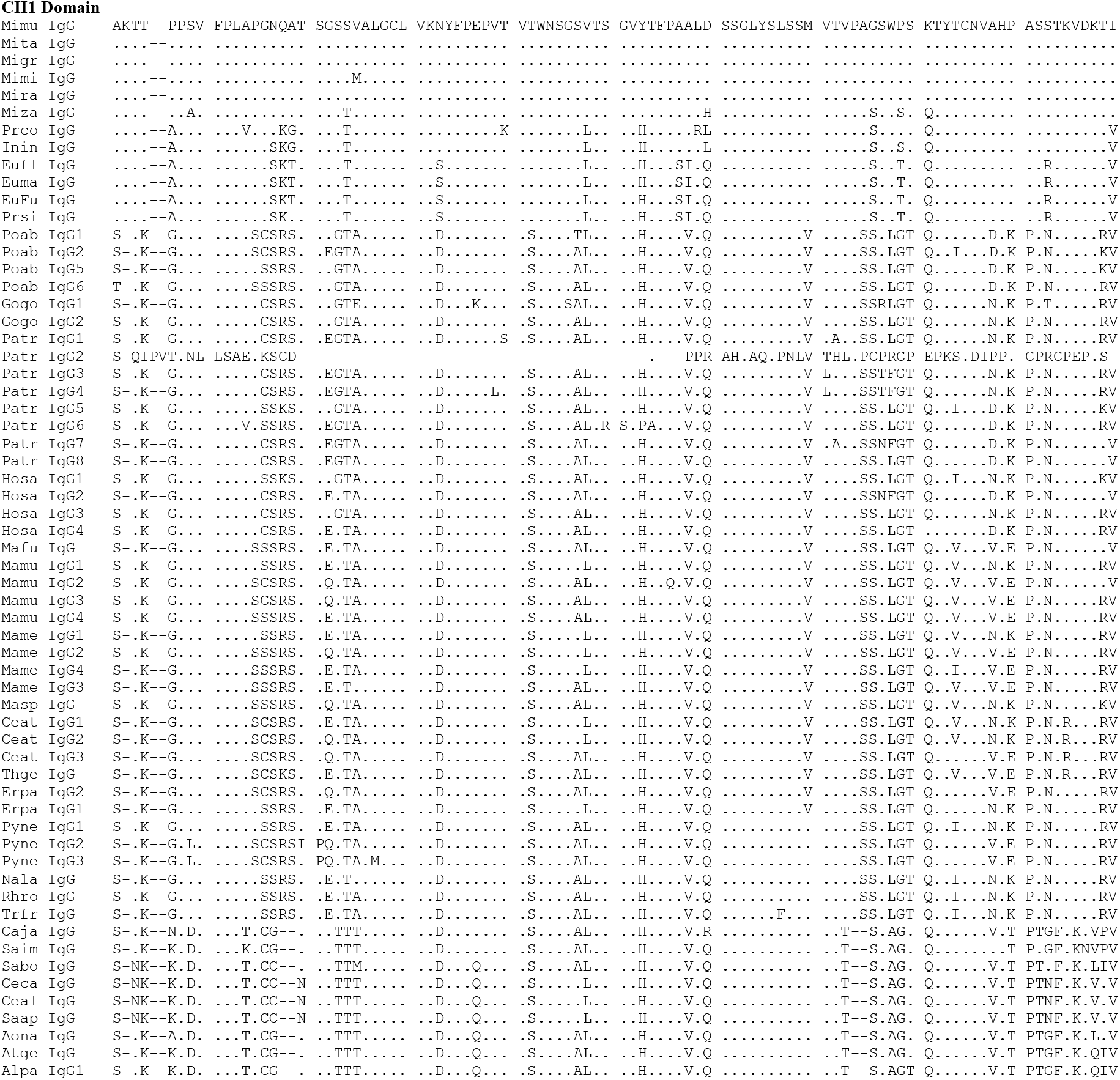
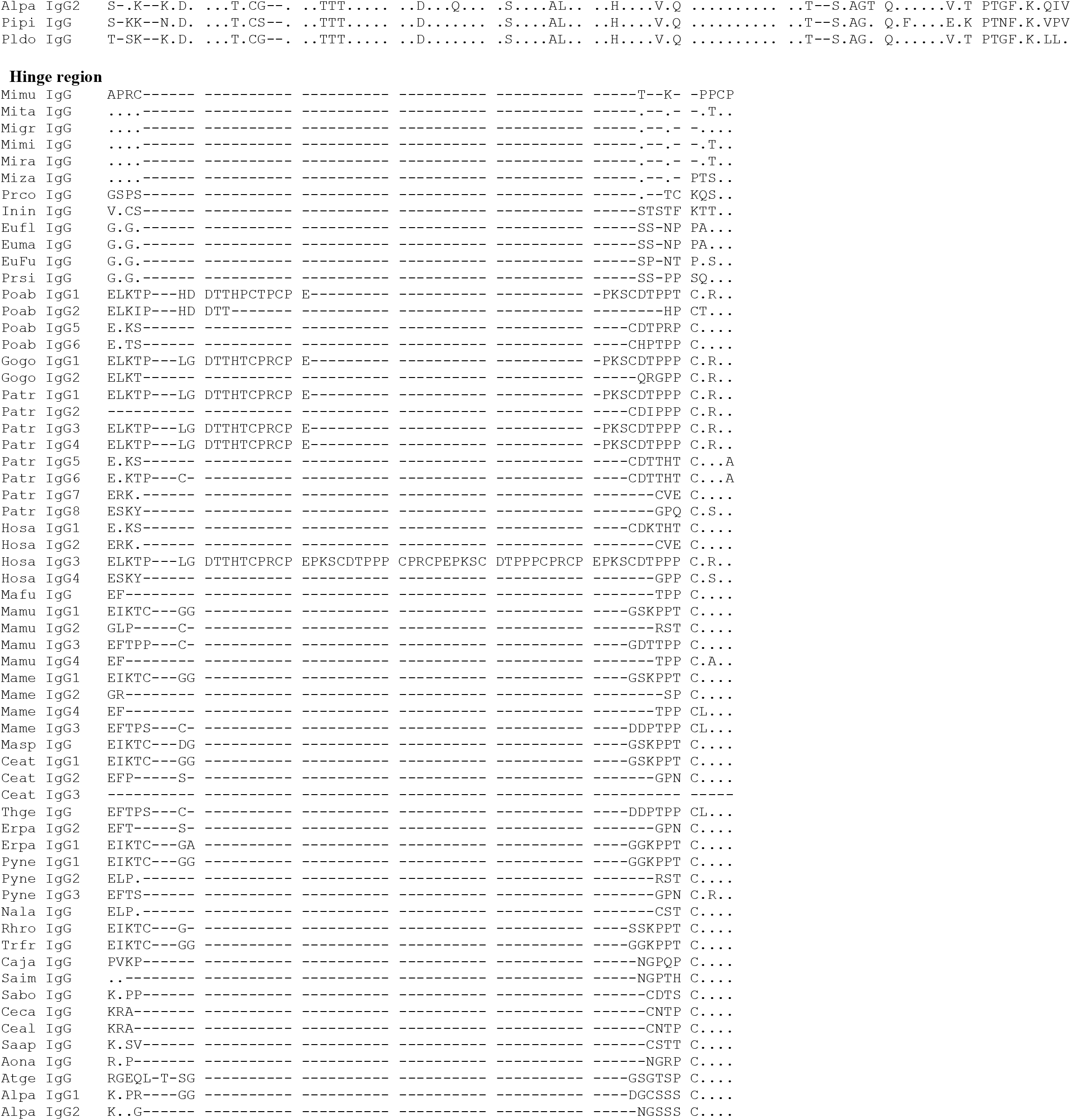
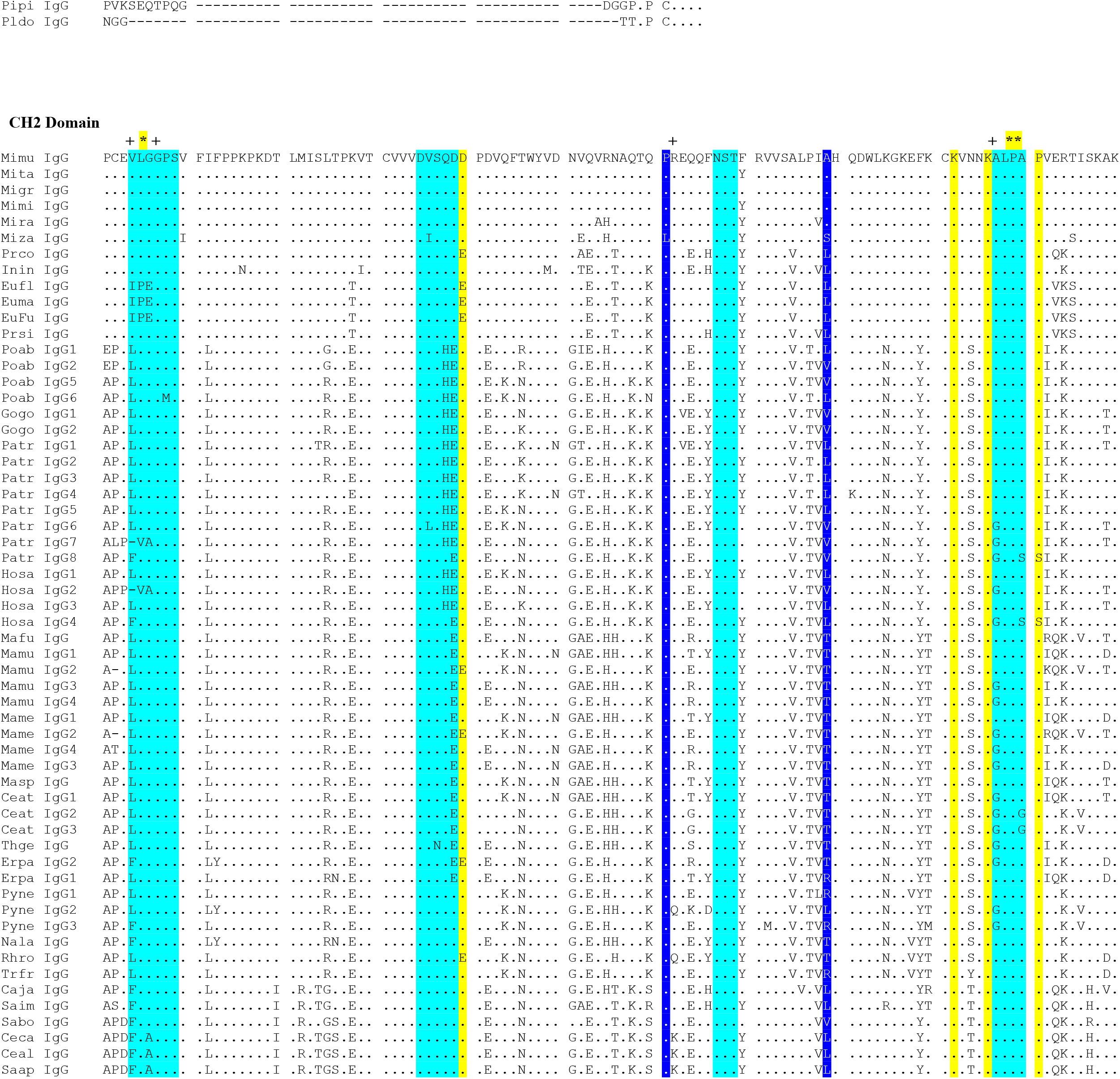
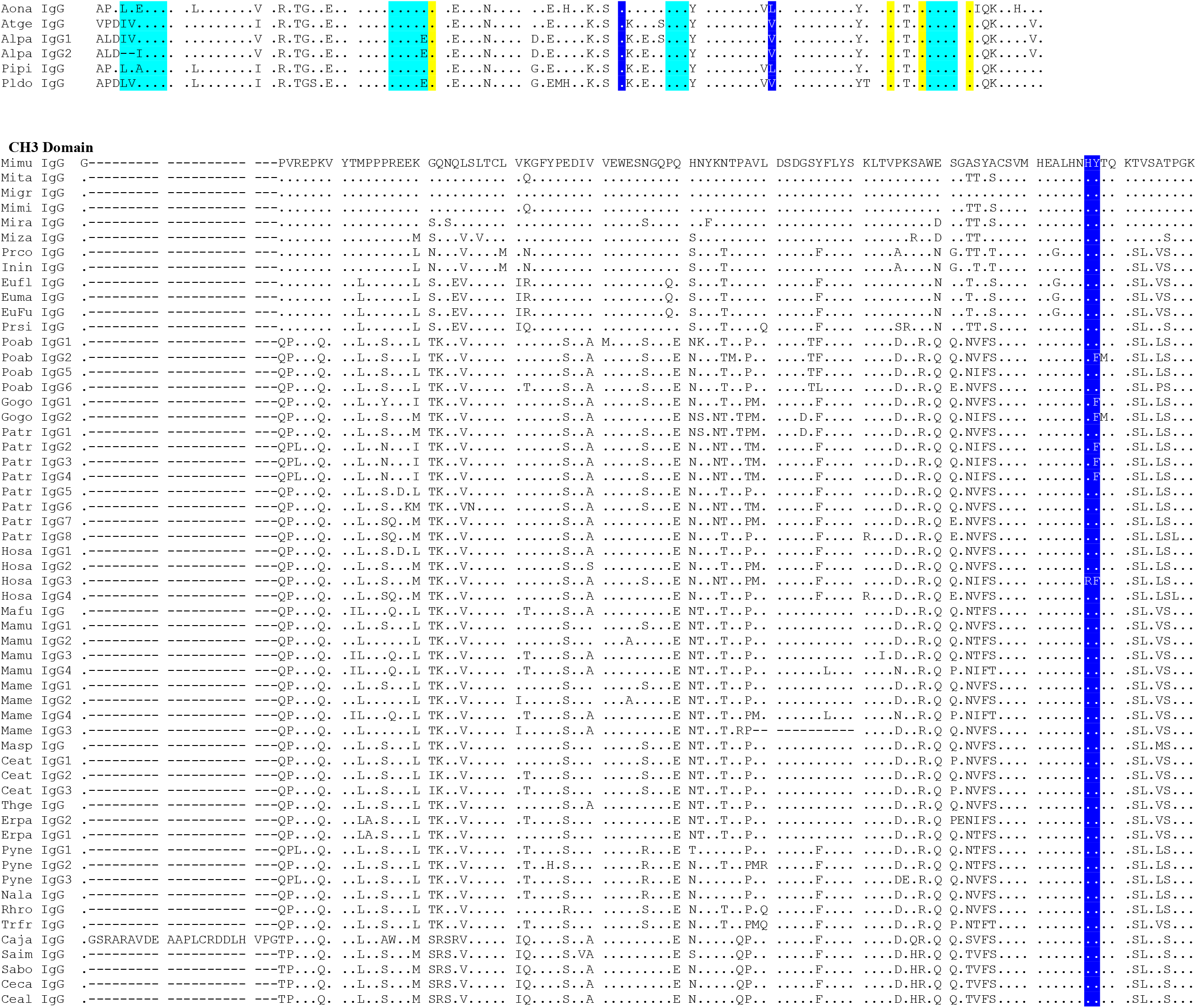
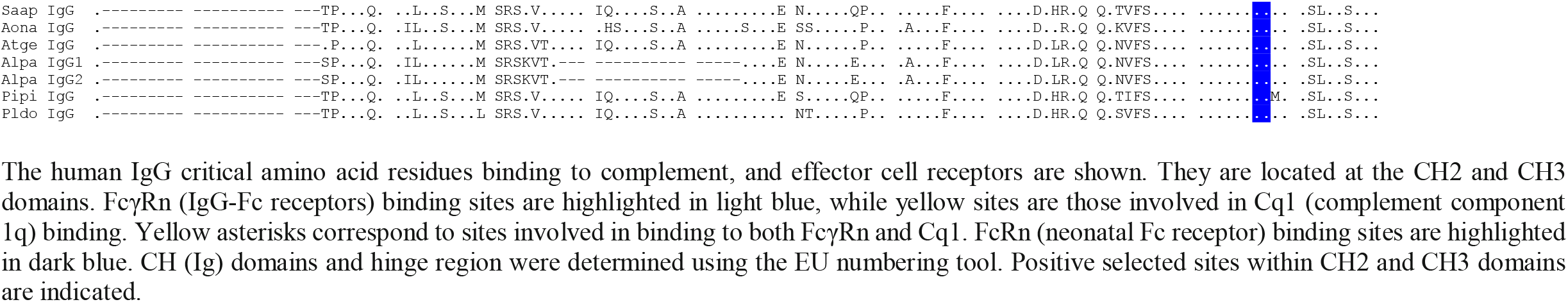
Alignment of deduced IgG amino acid sequences.

**Supplementary material 7.**
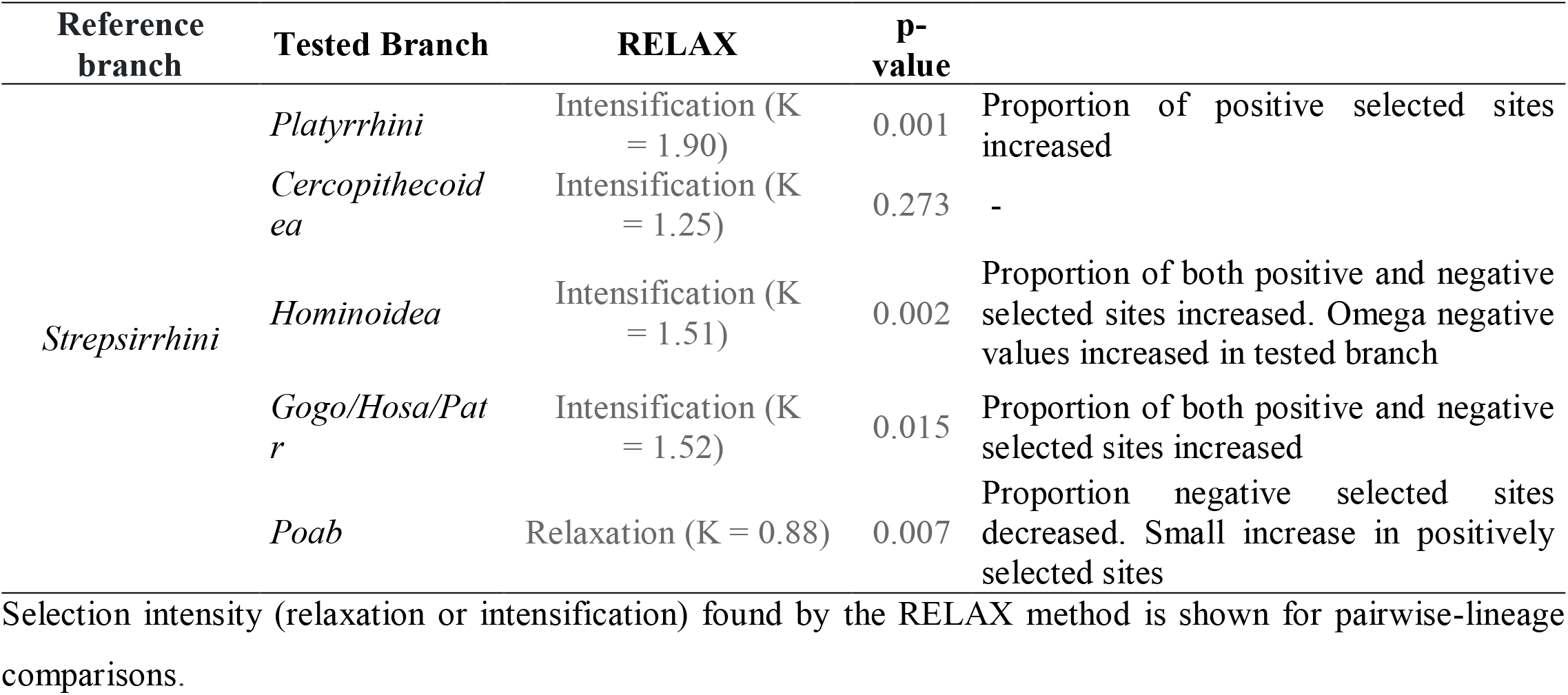
Relaxed or intensified natural selection between reference and tested branches.

